# Histone H3.3 lysine 9 and 27 control repressive chromatin states at cryptic *cis*-regulatory elements and bivalent promoters in mouse embryonic stem cells

**DOI:** 10.1101/2023.05.08.539859

**Authors:** Matteo Trovato, Daria Bunina, Umut Yildiz, Nadine Fernandez-Novel Marx, Michael Uckelmann, Vita Levina, Yekaterina Kori, Ana Janeva, Benjamin A. Garcia, Chen Davidovich, Judith B. Zaugg, Kyung-Min Noh

## Abstract

Histone modifications are associated with distinct transcriptional states, but it is unclear whether they instruct gene expression. To investigate this, we mutated histone H3.3 K9 and K27 residues in mouse embryonic stem cells (mESCs). Here, we find that H3.3K9 is essential for controlling specific distal intergenic regions and for proper H3K27me3 deposition at promoters. The H3.3K9A mutation resulted in decreased H3K9me3 at regions encompassing endogenous retroviruses and induced a gain of H3K27ac and nascent transcription. These changes in the chromatin environment unleashed cryptic enhancers, resulting in the activation of distinctive transcriptional programs and culminating in protein expression normally restricted to specialized immune cell types. The H3.3K27A mutant disrupted deposition and spreading of the repressive H3K27me3 mark, particularly impacting bivalent genes with higher basal level of H3.3 at promoters. Therefore, H3.3K9 and K27 crucially orchestrate repressive chromatin states at *cis*-regulatory elements and bivalent promoters, respectively, and instruct proper transcription in mESCs.

## Introduction

Specific histone PTMs are associated with gene expression through active or repressive chromatin states. For instance, histone H3 lysine 9 and lysine 27 acetylation (H3K9ac and H3K27ac) mark active *cis*-regulatory elements, whereas tri-methylation of these residues is a repressive mark, found at silenced repetitive elements (H3K9me3) and gene promoters (H3K27me3) [1–3]. Although the strong association between these PTMs and gene expression have been established [4, 5], a systemic study to investigate the precise contribution of specific histone residues and associated PTMs to the regulation of chromatin environments and gene expression is still lacking in mammals.

Histone H3 is present as three variants, H3.1-3. Canonical histone H3 (H3.1/H3.2) is produced from multiple gene copies and incorporated into nucleosomes upon DNA synthesis, thus being evenly distributed across the genome. In contrast, the incorporation of histone H3.3 is replication-independent. H3.3 differs from H3.1 and H3.2 by 5 and 4 amino acids, respectively [6]. The H3.3 specific residues are recognized by H3.3-specific chaperones, which deposit H3.3 at specific genomic regions, including actively transcribed genes and *cis*-regulatory elements, during interphase [7–11]. Histone H3.3 represents only ∼20% of the total H3 pool in proliferating cells [12] however, given that H3.3 is enriched at promoters, enhancers, repeat elements, and active genes, mutations of H3.3 result in substantial epigenome changes in mammalian cells and contribute to the pathogenesis of cancers [13–19]. Histone H3.3 is encoded by two genes (*H3f3a* and *H3f3b*), not located within the main histone gene clusters. The *H3f3a* and *H3f3b* double-knockout (KO) is embryonic lethal in mice, while single KOs are viable [20–22].

In this study, we investigated the role of H3.3K9 and H3.3K27 in transcriptional regulation and chromatin response by conducting histone H3.3 gene mutagenesis in mESCs [18, 19]. Loss of function lysine-to-alanine mutations in either the K9 or K27 residue resulted in significant transcriptional perturbation in mutant mESCs, accompanied by proliferation and/or differentiation defects. We found that H3.3K9 plays a crucial role in preserving specific heterochromatic regions and regulating cryptic cis-regulatory elements (CREs) derived from endogenous retroviruses (ERVs). An mESC-specific gene regulatory network revealed a connection between activated ERV-derived CREs and immune genes, as well as a subset of bivalent genes, that were selectively upregulated in H3.3K9A mutant cells. In contrast, H3.3K27 is directly required for transcriptional repression of a distinct subset of bivalent genes as their promoters were enriched with H3.3 and H3K27me3. These results provide evidence for an instructive role of H3.3 K9 and K27 residues in the regulation of gene expression. Our study also provides mechanistic insights into how H3.3K9 contributes to maintaining defined heterochromatic regions and controlling the expression of associated genes.

## Results

### H3.3K9A and K27A perturb transcription in mESCs

To elucidate the function of H3.3K9 and H3.3K27 and their modifications, we substituted K9 or K27 for alanine (H3f3a^−/−^; H3f3b^K9A/K9A^ or H3f3a^−/−^; H3f3b^K27A/K27A^) in mESCs and compared them to control mESCs (H3f3a^−/−^; H3f3b^WT/WT^) (**Fig. 1a** and **Fig. S1**). Hereafter, we refer to these lines as K9A, K27A, and control mESCs. After validating the chromosomal integrity of the edited lines (**Fig. S1f**), we assessed changes in the global levels of histone modifications and gene expression in three biological replicates of each mutant.

**Figure 1:**
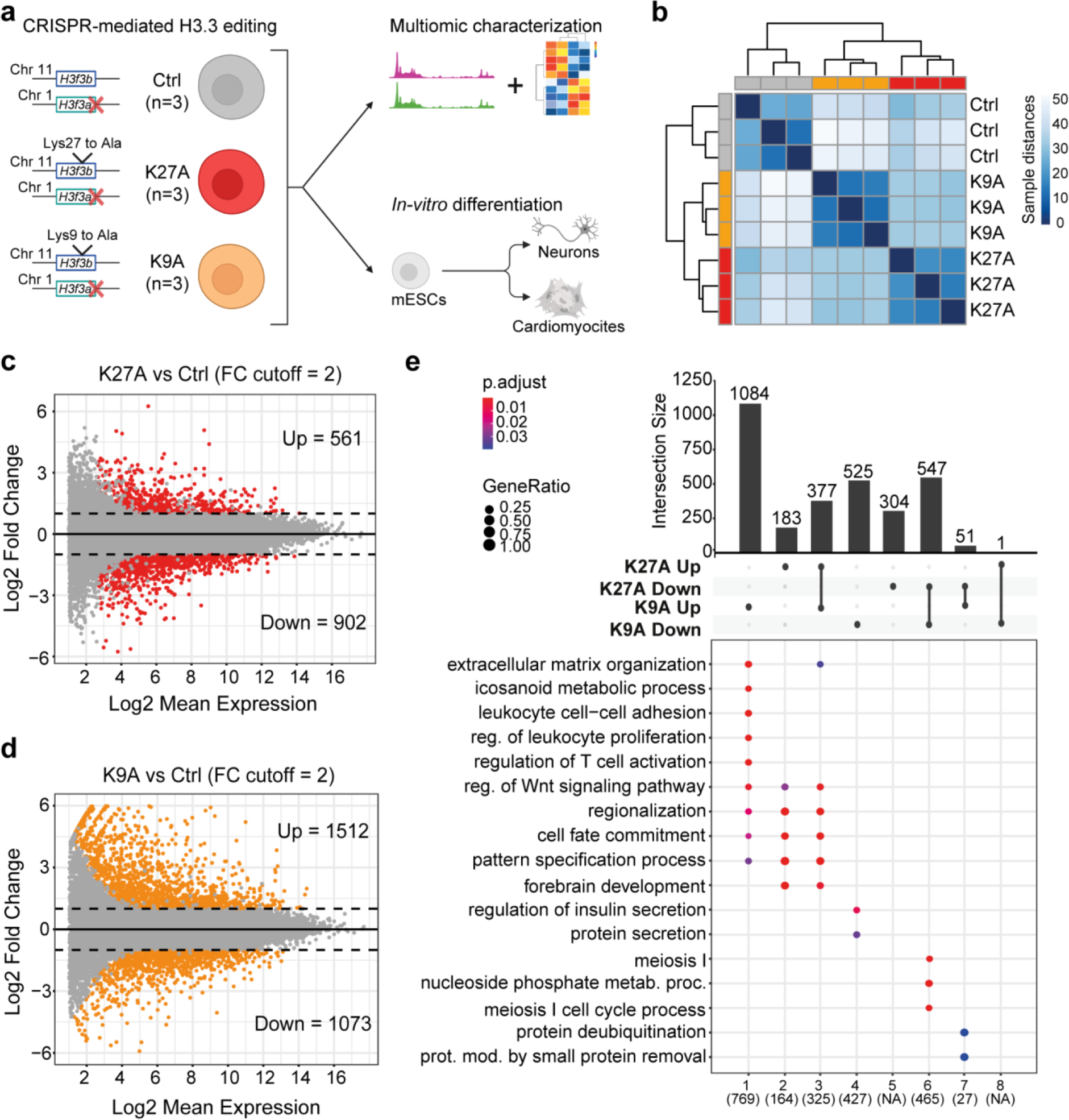
Transcriptional changes in H3.3 K27A and K9A mESCs. **(a)** Graphical outline with mESCs lines genotype. **(b)** Hierarchical clustering with sample-to-sample distances computed from rlog-transformed mRNA-seq counts. **(c)** MA plots of DEGs in K27A versus control mESCs and **(d)** DEGs in K9A vs. control mESCs. Colored genes represent significant DEGs, defined with padj<0.05 and log2(fold-change) cut-off of 1. **(e)** Upset plot of DEGs up-/down-regulated exclusively in one mutant or shared between the two mutants. On the bottom, the most significant GO terms for each group of genes are reported.

Histone Mass Spectrometry (MS) of the control and mutant lines confirmed the K9A and K27A substitutions in H3.3 proteins (**Fig. S2a**). MS also revealed the effects of the H3.3 mutations on global histone modification levels in H3.3 and H3.1/H3.2. For instance, we observed increased H3.1/H3.2K27me1 levels but unaltered H3.1/H3.2K27me2/3 levels in K27A mESCs compared to the control, indicating no compensatory effect on H3.1/H3.2K27me3 upon H3.3K27A substitution (**Fig. S2b**). In contrast, K9A mESCs showed decreased H3.3K27me1/2/3 levels and a reduction in H3.1/H3.2K27me1/2/3 levels as well as increases in histone H3.1/H3.2/H3.3 (H3K9ac, H3K14ac, H3K18ac, H3K23ac) and H4 acetylation (**Fig. S2a-c**), indicating broad changes in chromatin modifications upon H3.3K9 substitution.

To assess changes in gene expression, we performed mRNA-seq. Hierarchical clustering of the normalized read counts showed distinct transcriptional profiles for K9A and K27A cells, compared to controls (**Fig. 1b**). Analysis of differentially expressed genes (DEGs) revealed substantial changes for K9A (2585 DEGs, padj<0.05 & FC cut-off=2) and K27A mESCs (1463 DEGs, padj<0.05 & FC cut-off=2), with the number of DEGs and the magnitude of gene expression changes being greater in K9A than K27A (**Fig. 1c-d**).

To further characterize the gene expression changes in each mutant, we performed gene ontology (GO) enrichment analysis on the DEGs (**Fig. 1e**). Upregulated genes in K9A (n=1084) were related to immune response, extracellular matrix organization, and differentiation. Upregulated genes in K27A (n=183) and in both K9A and K27A mutants (n=377) were commonly enriched for development and differentiation processes, such as regionalization, pattern specification processes, and embryonic organ development. Downregulated genes in K9A were related to protein secretion processes (n=525) and, for both mutants, meiosis (n=547). Only 52 genes were differentially expressed in opposite directions in K9A and K27A, and these were enriched for protein removal processes. Thus, although H3.3 represents only ∼20% of the total histone H3 pool in mESCs, these single amino-acid substitutions lead to extensive changes in mESC transcription, indicating that H3.3 K9 and K27 are required for maintaining transcription in mESCs.

### H3.3K9A and K27A mESCs impaired differentiation

Timing of lineage-specific gene expression is critical during development, with chromatin modifiers and epigenetic modifications modulating cell fate transitions [23]. To understand the impact of H3.3K9A and H3.3K27A on developmental gene expression, we examined changes in marker gene transcription of embryonic lineages. Ectoderm and mesoderm marker genes were particularly affected in K9A and K27A mESCs, with lineage markers of trophectoderm (Cdx1, Cdx2), mesoderm (Tbxt, Snai1, Twist) and cardiac-mesoderm (Mef2c, Tbx1, Fgf10) showing significant upregulation (**Fig. 2a**).

**Figure 2:**
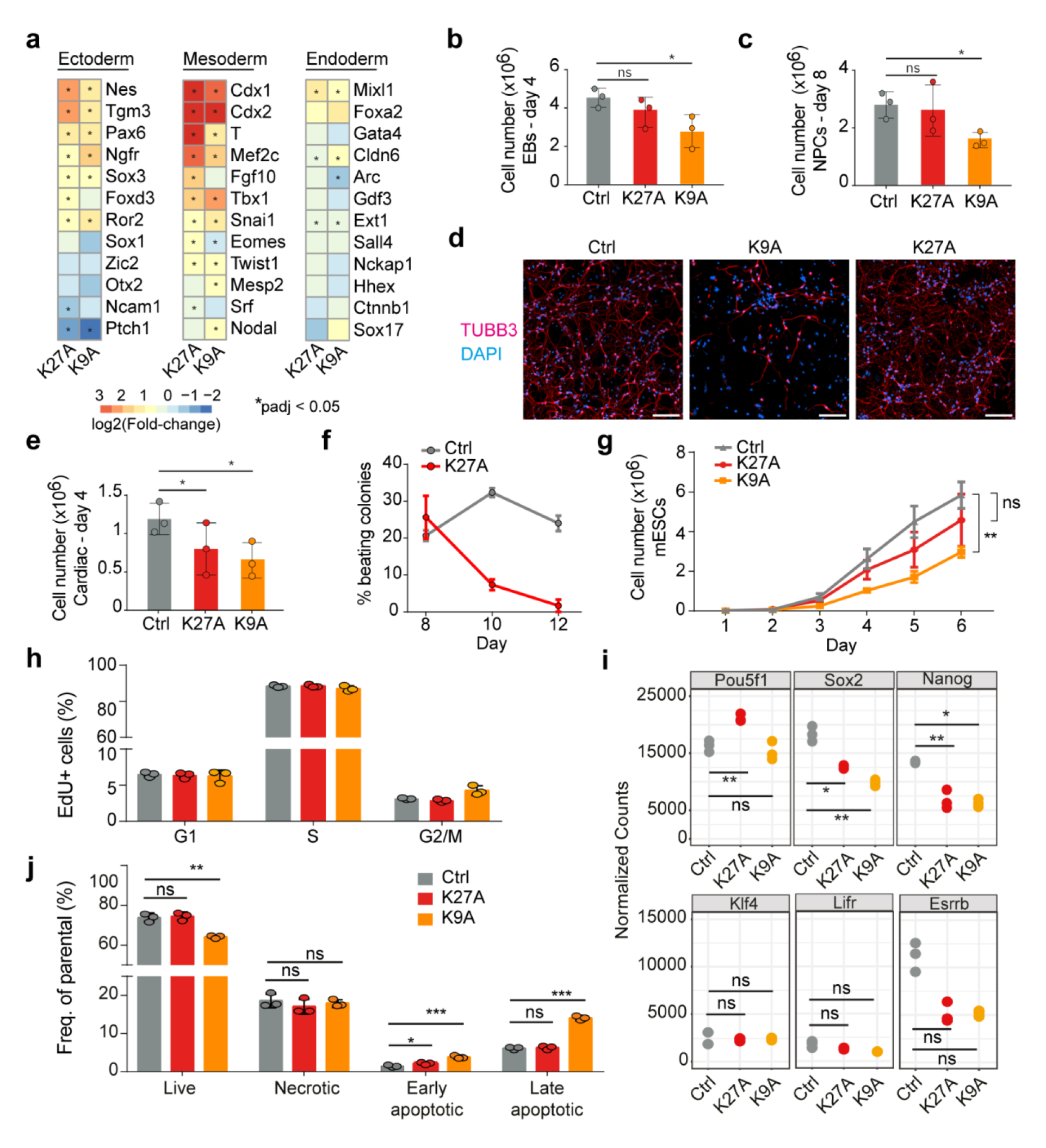
H3.3K9A mESCs fail to differentiate and display increased cell death. **(a)** Heatmap of log2(fold-change) expression for selected ectoderm, mesoderm, and endoderm marker genes in K27A and K9A mESCs; asterisks indicate whether the gene is significantly differentially expressed in the mutant compared to the control. **(b)** Barplot of cell numbers counted after dissociation of the embryoid bodies at day 4 and **(c)** day 8 of the *in-vitro* neuronal differentiation. **(d)** Merged immunofluorescence images of neurons on day 12 of the *in-vitro* differentiation, stained with antibodies against TUBB3 to detect neurons and with DAPI to detect nuclei. Scale bar 100 μm. **(e)** Barplot of cell numbers counted after dissociation of embryoid bodies at day 4 of *in-vitro* mesoderm-cardiac differentiation. **(f)** Line plot depicting the percentage of contracting colonies counted every two days throughout the final stage of the cardiac differentiation. **(g)** Growth curve for control and mutant mESCs. **(h)** Barplot with percentages of mESCs in G1, S, or G2/M phases detected following 5- ethynyl-2’-deoxyuridine (EdU) incorporation. **(i)** Normalized mRNA-seq counts for six selected pluripotency markers (Pou5f1/Oct4; Sox2; Nanog; Klf4; Lifr and Esrrb) in control, K27A and K9A mESCs. Significant differences were calculated with DESeq2 and adjusted with Benjamini Hochberg’s correction. (*:padj<0.05; **:padj<0.01; ns: padj>0.05). **(j)** Barplot with percentages of cells identified as live, necrotic, early apoptotic, or late apoptotic after Annexin V staining. In panels, **b**, **c**, **e-h**, and **j**, error bars represent the standard deviation of n=3 biological replicates. *P* value: unpaired t-test (*:p-value<0.05; **:p-value<0.01; ***:p-value<0.001; ns:p-value>0.05).

We investigated whether the activation of such genes in mutant mESCs could affect *in-vitro* differentiation. Upon differentiation, K9A mESCs formed significantly smaller embryoid bodies (EBs) (**Fig. 2b**), and upon subsequent induction to neuro-ectoderm, they generated fewer neuronal progenitor cells (NPCs), failed to differentiate into neurons, and formed a heterogeneous population of cells characterized by a non-neuronal-like morphology, compared to control mESCs (**Fig. 2c-d**). K27A cells formed a homogeneous population of neurons with regular morphology, unlike K9A cells. NPCs marker expression (Hash1, Neurod1, Pax6, and Sox2) at day 8 of differentiation was significantly affected only in the H3.3K9A mutant (**Fig. S3a**). Immunofluorescence staining for neuronal markers (Map2 and beta-III tubulin) at day 12 of differentiation showed a marked decrease in K9A but not in K27A cells (**Fig. 2d** and **Fig. S3b**).

Cardiac-mesoderm differentiation of K9A mESCs revealed a significantly reduced number of cells at the intermediate stage of the differentiation (**Fig. 2e**). The K9A cardiac-mesoderm EBs failed to progress further differentiation. Cardiac-mesoderm EBs derived from K27A mESCs also exhibited reduced cell numbers at day 4 of differentiation (**Fig. 2e**) but were able to commit to the mesoderm lineage. However, the number of actively contracting colonies gradually declined during differentiation (**Fig. 2f**), indicating that K27A cells are unable to maintain a functional cardiomyocyte differentiated state.

### Growth delay and cell death in H3.3K9A mESCs

The differentiation defects in H3.3K9A mESCs imply that factors that regulate differentiation have been perturbed. Monitoring proliferation over six days revealed decreased numbers of K9A mESCs, but not of K27A mESCs, compared to the control (**Fig. 2g**). This could be due either to reduced proliferation and/or increased cell death. Cell cycle analysis in controls and mutants revealed the same percentage of cells in G1, S, or G2/M phases (**Fig. 2h** and **Fig. S3c**), indicating that proliferation was unaffected in K9A and K27A mESCs. Pluripotency marker gene expression was overall maintained, except for Nanog and Sox2, which were decreased in both K9A and K27A mESCs (**Fig. 2i**). However, annexin V staining revealed significantly increased cell death in K9A mESCs, but not in K27A mESCs (**Fig. 2j** and **Fig. S3d**), which was uncoupled to the accumulation of lipid reactive oxygen species (**Fig. S3e**). Altogether, these results indicate that increased cell death causes reduced cell numbers in K9A mESCs.

### Distal regulatory element activation in H3.3K9A mESCs

H3.3 K9A and K27A mESCs are viable, able to self-renew, and display substantial changes in gene expression compared to wild type. Therefore, using ChIP-seq, we explored the impact of H3.3 substitutions on the chromatin environment in mESCs. H3K9me3 ChIP-seq and differential analyses revealed that, among ∼48300 H3K9me3 regions, 8685 were differentially abundant (DA) in K9A mESCs, while only 2 were changed in K27A mESCs (**Fig. 3a** and **Fig. S4a**). 8203 out of 8685 DA-H3K9me3 regions in K9A mESCs showed a reduced signal, compared to the control and the majority of them were located within heterochromatin (**Fig. 3a**). Genomic annotation of the DA-H3K9me3 regions revealed that: 68% were located in distal intergenic regions, 18% were in introns, 3% were within 2 kb of a transcriptional start site (TSS), and the remainder were located in exons and 5’-/3’-UTRs (**Fig. S4b**). ChIP-seq of H3K27ac in control and K9A mESCs revealed a significant increase of this activating mark in 13177 regions (**Fig. 3b**), mostly in regions annotated as heterochromatin (**Fig. 3b**), overlapping with the H3K9me3 decreased regions (**Fig. S4b**).

**Figure 3:**
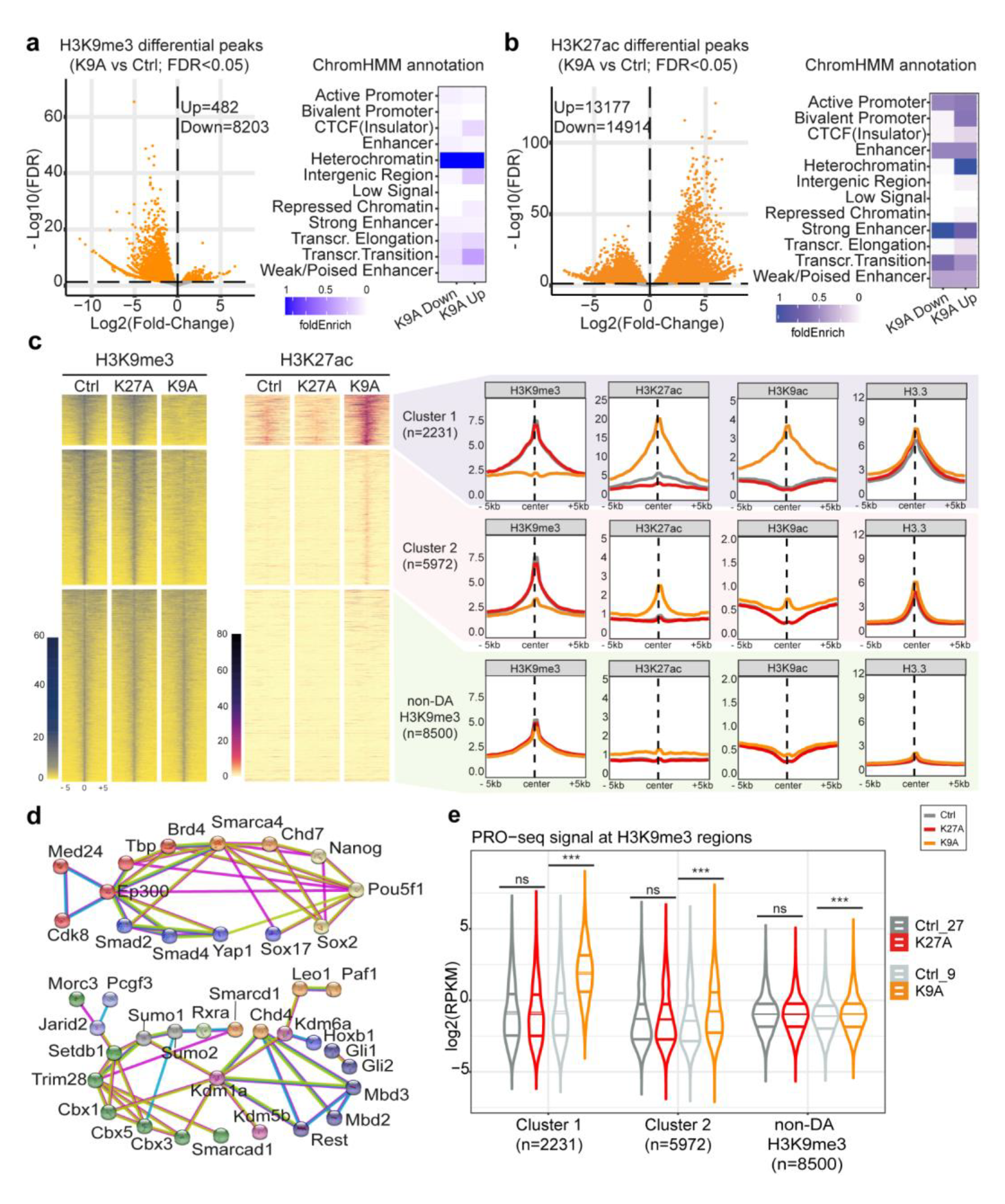
H3K9me3 reduction with gaining in H3K27ac and transcription at distinct heterochromatic regions in H3.3K9A mESCs. **(a)** Volcano plot showing differential H3K9me3 ChIP-seq peaks in K9A vs. control mESCs (consensus peakset with n=48346 broad peaks) and ChromHMM annotation of DA-H3K9me3 regions. **(b)** Volcano plot showing differential H3K27ac ChIP-seq peaks in K9A vs. control mESCs (consensus peakset with n=94000 peaks) and ChromHMM annotation of DA-H3K27ac regions. **(c)** Heatmaps of H3K9me3 and H3K27ac ChIP-seq signals at H3K9me3 regions (summit ± 5 kb). Differentially abundant regions are divided into two clusters (k-means clustering) and sorted from a high-to-low H3K9me3 signal in control. ChIP-seq signal calculated as the average of three replicates for control or mutant mESCs, in 250 bp sliding genomic windows. On the right, metaprofile plots with H3K9me3, H3K27ac, H3K9ac, and H3.3 average ChIP-seq signals are reported. **(d)** STRING network analysis of chromatin and transcription factors enriched at Cluster 1 (top) and Cluster 2 (bottom) DA-H3K9me3 regions. Colors indicate the chromatin complex/category (red: transcriptional co-activator; orange: chromatin remodeler/elongation complex; yellow: pluripotency TF; blue: developmental TF; purple: histone demethylase; green: heterochromatin factor; dark-blue: transcriptional repressor; light-blue: Polycomb subunit). STRING settings: physical subnetwork (the edges indicate that the proteins are part of a physical complex) – medium confidence (interaction score 0.4). **(e)** Nascent transcription signal at DA-H3K9me3 regions reported as log2(RPKM) values. *P* value: unpaired t-test (***: p-value<0.001; ns: p-value>0.05). PRO-seq experiments were performed independently for K27A and K9A. Control samples were repeated in each batch and reported in two different gray shades.

To better understand the de-repressed H3K9me3 regions, we employed k-means clustering to group the DA-H3K9me3 peaks (n=8203) based on their H3K9me3 signal. This resulted in two distinct clusters: Cluster 1, which fully lost the H3K9me3 signal in the K9A mutant, and Cluster 2, where the signal was reduced yet still present. For comparison, we included 8500 randomly selected non-DA H3K9me3 peaks in the analysis as a control set (**Fig. 3c**). Compared to the non-DA regions, both Cluster 1 and 2 displayed elevated H3.3 deposition, indicating that H3.3 is the primary contributor to the H3K9me3 signal in these regions. Cluster 1 regions, compared to Cluster 2, showed a higher basal level of H3K27ac in control mESCs, which was further increased in K9A mESCs. The H3K9ac signal was increased in K9A mESCs, despite the enrichment of mutant H3.3, indicating the presence of H3.1/H3.2 in the H3K9ac-enriched regions. Cluster 2 regions contained no basal histone H3 acetylation, regardless H3K9me3 levels were reduced, and there was a modest increase in H3K27ac and H3K9ac in K9A mESCs (**Fig. 3c**).

Given that Cluster 1 and 2 regions exhibited differences in H3K27ac upon H3.3K9A substitution, we used the ChIP-Atlas database [24] to compare the occupancy of chromatin modifiers and transcription factor (TF) binding between these two sets of regions in wild-type mESC. Consistent with higher basal levels of H3K27ac, Cluster 1, compared to Cluster 2, showed a relative enrichment of transcriptional co-activators (P300, BRD4, MED24), subunits of chromatin remodelers (SMARCA4, CHD7), and TFs linked to pluripotency (NANOG, SOX2, OCT4/POU5F1) and development (SOX17, SMAD2/4, YAP1). Cluster 2 showed a relatively higher enrichment for core heterochromatin factors, such as HP1 proteins (CBX1, CBX3, CBX5), KAP1/TRIM28, SETDB1, and other transcriptional repressors (the STRING network of the factors shown in **Fig. 3d**).

To determine if these de-repressed H3K9me3 regions elicit transcriptional activity, we assessed nascent transcript levels (PRO-seq). Cluster 1 regions, which displayed a larger H3K9me3 loss and H3K27ac increase, showed a marked gain in nascent transcription (**Fig. 3e**). Cluster 2 regions showed a more modest increase in nascent transcription, consistent with the ChIP-seq data. Altogether, our results indicate that the H3.3K9 residue contributes to maintaining a set of distal intergenic regions in a repressed state and that the H3K9me3 signal reduction initiates a switch to an active chromatin state.

### De-repressed cryptic enhancers in H3.3K9A mESCs upregulate immune-related genes

We hypothesized that the de-repressed distal elements in Cluster 1 and 2 could act as cryptic enhancers, driving the expression of upregulated genes in K9A mESCs. To test this, we examined the expression of genes within 50 kb of the DA-H3K9me3. Genes close to DA-H3K9me3 regions (especially Cluster 1 regions) were, on average, selectively upregulated in H3.3K9A mESCs (**Fig. S4d**). To further refine the putative enhancer-gene connections, we constructed an enhancer-mediated gene regulatory network (GRN) using GRaNIE [25], based on the co-variation between mRNA-seq and H3K27ac ChIP-seq data, from a total of 12 mESCs samples (see Methods). The GRN allows us to infer interactions between *cis*- regulatory elements (i.e., H3K27ac peaks) and genes. After filtering for significant connections, we obtained 16701 CRE-gene connections, composed of a network of 5229 genes and 5694 regulatory elements. In the GRN, a total of 657 DA-H3K9me3 regions were retained, which were mostly annotated as distal intergenic and intronic regions. These DA-H3K9me3 regions were linked to 789 genes (n=729 in Cluster 1 and n=118 in Cluster 2, with n=58 genes that were linked to both clusters; average peak-gene distance ≈ 108 kb) and 82% (n=651/789) of them were upregulated in K9A mESCs (**Fig. 4a**).

**Figure 4:**
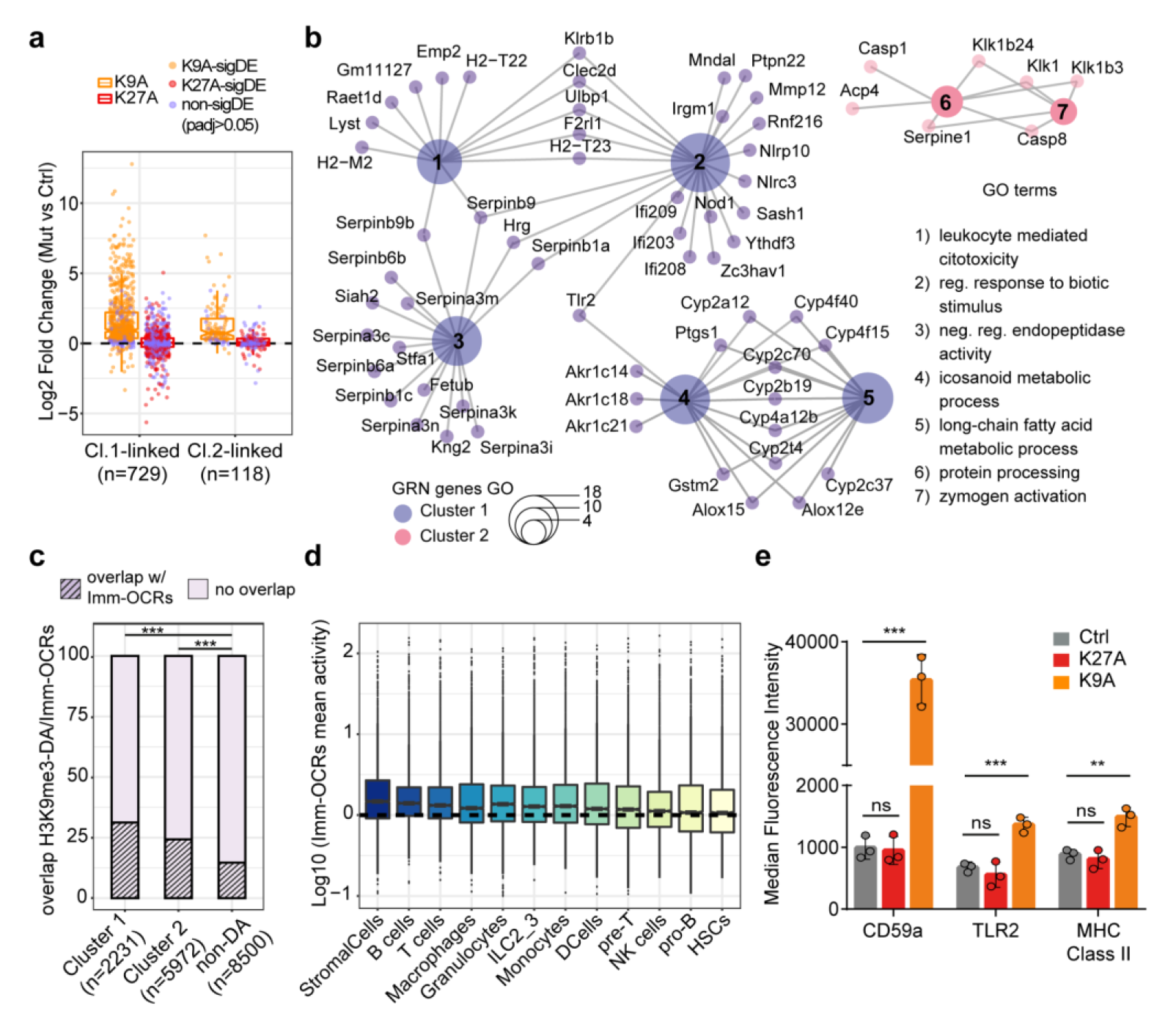
Cryptic CREs activated in H3.3K9A mESCs drive the expression of immune-related genes. **(a)** Expression of genes connected to DA-H3K9me3 regions through the GRN; log2(fold-change) values calculated with DESeq2 are plotted, and individual genes (data points) are colored to indicate if the differential expression is significant (orange/red) or not (purple). **(b)** Cnet plot of gene ontology terms showing enrichment for genes connected to Cluster 1 (purple) and 2 (pink) DA-H3K9me3 regions included in the GRN. **(c)** Stacked barplot displaying the percentage of overlap between DA-H3K9me3 regions and Imm-OCRs. *P* value: a two-proportions Z-test (***: p-value<0.001). **(d)** Mean activity of Imm-OCRs overlapping with DA-H3K9me3 regions in immune cell populations; cell types are sorted by decreasing the mean activity of the open chromatin regions. **(e)** Barplots of median fluorescence intensity of selected surface markers. Standard deviation from three biological replicates of control (gray), K27A (red), or K9A (orange) mESCs. *P* value: unpaired t-test (**: p-value<0.01; ***: p-value<0.001; ns: p-value>0.05).

Many genes specifically upregulated in K9A mESCs were related to immune processes (**Fig. 1e**). We tested if the upregulation of immune-related genes could be driven by newly activated cryptic CREs. GO analysis of the genes connected to de-repressed CREs via the GRN revealed terms such as leukocyte-mediated cytotoxicity and regulation of response to a biotic stimulus (**Fig. 4b**). Individual genes underlying these terms included Toll-like receptor genes (Tlr2), class II major histocompatibility complex genes (MHC, e.g., H2-M2 and H2- T23), interferon-inducible genes (Mndal), and members of the nucleotide-binding leucine-rich repeat gene family (Nod1) (**Fig. 4b**). Two additional GO terms related to inhibition of protease activities, including members of the serine protease inhibitors (serpins), and fatty acid and eicosanoid metabolism, both with reported roles in innate immunity [26, 27], also showed enrichment among the upregulated genes linked to the de-repressed CREs in K9A mESCs.

To test whether the H3K9me3-marked cryptic CREs are active in immune cell types, we retrieved data from the mouse immune-system *cis*-regulatory atlas, which comprises over 500,000 putative CREs from 86 primary immune cell types, identified using ATAC-seq [28]. The comparison between the DA-H3K9me3 regions and the collection of immune open-chromatin regions (Imm-OCRs) revealed a significant amount of overlap in Cluster 1 (31%; n=697/2231) and Cluster 2 (24%; n=1450/5972), compared to the overlap in non-DA H3K9me3 regions (14%; n=1247/8500) (**Fig. 4c**). To determine if these CREs from Cluster 1 and 2 (n=2147) are active in specific immune cells, we merged highly correlated Imm-OCRs from closely-related cell types and sorted the resulting cell types based on the activity (i.e., chromatin openness) of overlapping Imm-OCRs (**Fig. 4d**). Apart from stromal cells, the highest activity was observed in specialized immune cell types such as B-cells, T-cells, macrophages, and granulocytes. This result indicates that a significant fraction of the cryptic CREs de-repressed in H3.3K9A mESCs function as enhancers active in specialized immune cell types.

To examine whether immune-related TFs are active in H3.3K9A mESCs, we performed a differential TF activity analysis using diffTF [29]. By comparing K9A and control mESCs (padj<0.001), we identified 84 differentially active TFs. HIVEP1/ZEP1 and HIVEP2/ZEP2, which bind viral promoters and CREs of immune-related genes [30, 31], were amongst the TFs with the highest activity in K9A mESCs (**Fig. S5a**). Other top hits included three NF-kB subunits (REL/C-REL; RELA/p65; NFKB1/p50), with roles in inflammation and in the immune response [32, 33], and TAL1, a TF involved in hematopoiesis [34, 35].

To determine whether transcriptional upregulation of immune genes in K9A mESCs translated into protein expression, we tested three targets: CD59a (a surface glycoprotein involved in T-cell activation and inhibition of the complement membrane attack complex), TLR2 (a surface protein required for activation of innate immunity), and MHC Class II (surface proteins typically expressed in immune cells dedicated to the antigen presentation). Immunolabeling followed by flow cytometry analysis revealed that all three surface proteins were significantly more expressed in K9A mESCs, compared to control mESCs, whereas their expression was unchanged in K27A mESCs (**Fig. 4e**). Thus, the de-repressed CREs in H3.3K9A mESCs regulate the expression of immune-related genes, culminating in the production of proteins restricted to specialized cell types.

### Cryptic enhancers in H3.3K9A mESCs derive from ERVs

Having identified H3.3-marked genomic regions acting as CREs upon the reduction of H3K9me3 level, we sought to elucidate the unique features of these loci. Histone H3.3 is enriched at ERVs [36–38], which are a repertoire of genomic sequences with regulatory potential [39–42] and represent 15-30% of CREs in immune cells [43–46]. We investigated whether the cryptic CREs are derived from ERV sequences by assessing the degree of overlap between de-repressed CREs in K9A mESCs and ERVs. Analysis revealed that 60% (n=1347/2231) of the DA-H3K9me3 regions in Cluster 1 and 40% (n=2322/5972) in Cluster 2 displayed overlap with ERVs (**Fig. 5a**), which was significantly higher than 22% (n=1916/8500) in the control set of non-DA H3K9me3 regions (**Fig. 5a**). The high presence of ERVs in these H3K9me3-marked genomic regions indicates their inherent potential to act as CREs.

**Figure 5:**
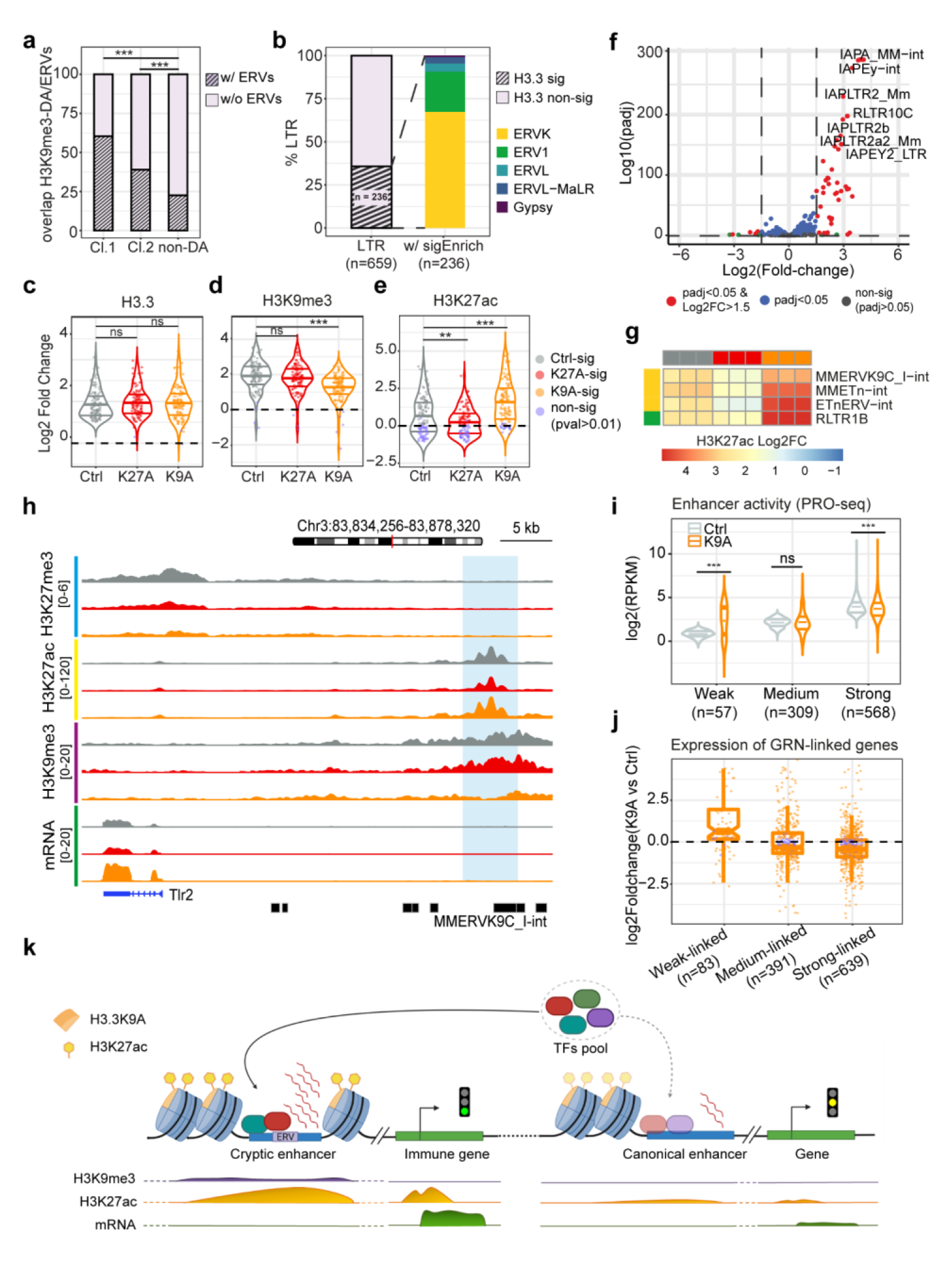
Cryptic CREs activated in H3.3K9A mESCs derive from ERVs. **(a)** Stacked barplot showing the percentage of DA-H3K9me3 regions with at least one ERV. *P* value: a two-proportions Z-test. **(b)** Stacked barplot indicating the percentage of LTR elements with significant (pval<0.01) H3.3 enrichment (left), and the classification of ERV families with significant H3.3 signal (right). **(c-e)** Violin plot displaying **(c)** H3.3, **(d)** H3K9me3, **(e)** H3K27ac log2(Fold-change) signal at ERVs subfamilies (n=108), in control, K27A and K9A mESCs. Individual ERV subfamilies (data points) are colored to indicate if the ChIP enrichment is significant (gray/red/orange) or not (purple). *P* value: unpaired t-test. **(f)** Volcano plot showing differential LTRs expression in K9A vs. control mESCs. **(g)** Heatmap displaying H3K27ac log2(Fold-change) enrichment at four selected ERV subfamilies (complete heatmap in Fig. S6g). **(h)** Genome browser snapshot of a representative Cluster 1 DA-H3K9me3 region (highlighted), linked through the GRN to the Tlr2 gene; H3K27me3, H3K27ac, H3K9me3, and mRNA-seq tracks are shown. For ERVs annotation, only the name of the ERV located within the DA-H3K9me3 region is shown. **(i)** PRO-seq signal at Weak, Medium, and Strong dREG enhancers included in the GRN for K9A and control mESCs. Log2(RPKM) values are reported. *P* value: unpaired t-test. **(j)** Expression of DEGs connected to dREG enhancers through the GRN; log2(fold-change) values calculated with DESeq2 are plotted, and individual genes (data points) are colored to indicate if the differential expression is significant (orange) or not (purple). **(k)** Model of *cis*-regulatory elements rewiring in H3.3K9A mESCs. *P* value in **a,c-e,i**: ***: p-value<0.001; **: p-value<0.01; ns: p-value>0.05. Data from three biological replicates of control, K9A, or K27A clonal lines.

To study the H3.3K9A-induced ERVs activation and their subfamilies comprehensively, we examined the ChIP-seq signals of H3.3, H3K9me3, and H3K27ac at repeat elements in control and mutant mESCs. Histone H3.3 was significantly enriched at 35% (n=236/659, pval<0.01) of ERV subfamilies, including ERVK, ERV1, and ERVL (**Fig. 5b**). Analyses of ERV subfamilies with higher H3.3 enrichment (n=108; log2 Fold-change cut-off=1) revealed that although H3.3 occupancy was unaffected in the mutants (**Fig. 5c**), H3K9me3 levels were significantly reduced in K9A mESCs, and accompanied by an increase in H3K27ac levels (**Fig. 5d-e**). H3K9me3 ChIP-qPCR confirmed the specific signal reduction at ERVs (**Fig. S6c**). Furthermore, mRNA-seq and RT-qPCR analyses of LTRs expression demonstrated an increase in ERVs transcriptional activities, selectively in K9A mESCs (**Fig 5f** and **Fig. S6d-f**), consistent with the ChIP-seq analyses.

Among the ERV subfamilies, RLTR1B, ETnERV-int, MMETn-int, and MMERVK9C I-int, which showed a modest basal H3K27ac signal in control mESCs (**Fig. S6g**), displayed particularly high gain of the H3K27ac signal in K9A mESCs (**Fig. 5g**), indicating their potential role as active cryptic CREs. Inspection of Cluster 1 and 2 regions revealed MMERVK9C I-int and MMETn-int within regions linked through the GRN to immune-related genes that were upregulated in K9A mESCs, such as Tlr2 and Cd59a, respectively (**Fig. 5h** and **Fig. S7a**). Collectively, these results indicate that a large fraction of the cryptic CREs activated in H3.3K9A mESCs is ERV-derived enhancers, and some of them are capable of driving the expression of immune-response genes.

Besides functioning as cryptic CREs, de-repression of ERVs and production of endogenous viral transcripts could trigger an interferon-mediated antiviral response, also known as viral mimicry [47, 48]. However, in mESCs, viral infection or exposure to synthetic viral RNA does not result in interferon responses due to low expression levels of major innate immunity factors [49, 50]. Consistent with this, we did not observe a global upregulation of interferon-stimulated genes in K9A mESCs (**Fig. S7b**), indicating that viral mimicry is not the main cause for the observed activation of immune genes.

### Enhancer rewiring in H3.3K9A mESCs

Given the de-repression of numerous ERV-derived CREs with regulatory potential in K9A mESCs, we predicted a change in the distribution of transcription factors and chromatin components involved in gene regulation. Analysis of the PRO-seq data with the dREG and tfTarget packages [51–53] revealed 98 and 20 TF motifs significantly enriched at CREs with increased and reduced activity in K9A mESCs, respectively. TF motifs at more active CREs were related to immune cell specification (Bhlhe40, Tcf3, Irf) and NF-kB pathway, as well as developmental processes (Tbx2, Tbx3, Sox9, Sox17, Hoxa3, Hoxd3) and maintenance of pluripotency (Pou5f1/Oct4, Sox2, Myc). TF motifs at CREs with reduced activity were related to cellular function (Fos, Jun) and pluripotency (Pou5f1/Oct4, Esrrb) (**Fig. S8a-b**), suggesting a decrease in the activity of pre-existing canonical enhancers. We also observed that besides the increase in 13177 H3K27ac peaks, 14914 H3K27ac peaks were reduced in K9A mESCs (**Fig. 3b**).

We investigated whether the activity of canonical enhancers is altered in K9A mESCs concomitantly with the activation of cryptic enhancers and whether this reflects changes in gene expression. Among the active canonical enhancers identified using dREG, 934 were linked to at least one gene in the GRN. We divided these into three groups based on their nascent transcript reads in control mESCs (n=568 strong; n=309 medium; n=57 weak). In K9A mESCs, nascent transcription levels were mainly reduced at strong enhancers, medium enhancers did not show a significant decrease, and weak enhancers showed increased activity (**Fig. 5e**), even though the H3K27ac signal was significantly reduced across all three groups (**Fig. S8c**). We identified 639, 391, and 83 DEGs connected to strong, medium, and weak enhancers, respectively (average peak-gene distance ≈ 111 kb). Consistent with the reduction in enhancer activity, genes connected to strong enhancers were, on average, downregulated (**Fig. 5f**). Genes connected to medium and weak enhancers either showed a trend towards downregulation or significant upregulation, respectively, correlating with the differences in enhancer activity.

As a comparison, given that H3K27ac showed a marked decrease also in K27A mESCs (**Fig. S8d**), we analyzed canonical enhancer activity and associated gene expression in K27A mESCs. Nascent transcription levels were significantly reduced at both strong and medium enhancers (**Fig. S8f**), correlating with overall reduced expression of connected genes (**Fig. S8g**). The correlation between enhancer activity and associated gene expression was not observed in a recent study with H3.3K27R mESCs [17], implying a stronger effect of the H3.3K27A substitution at CREs. Indeed, assessing H3K9ac and H3K18ac signals revealed their decreases in K27A mESCs (**Fig. S8e** and **Fig. S8h**), which was not seen in K27R mESCs [17] nor in K9A mESCs.

Taken together, these analyses indicate that substantial changes in the chromatin state of H3.3K9me3-marked CREs reshape gene regulatory networks in K9A mESCs, with de-repressed ERV-derived CREs upregulating a subset of genes in K9A mESCs, and potentially providing a plethora of TFs binding sites [39, 54], resulting in a redistribution of TFs and subsequent gene expression changes in K9A mESCs (**Fig. 5k**).

### H3K27me3 reduction at bivalent promoters in H3.3K9A mESCs

Having detailed immune-response genes upregulated in K9A mESCs, we studied genes associated with developmental processes, many of which were upregulated in both K9A and K27A mESCs (**Fig. 1e**). We focused on the promoter region of these developmental genes, which are typically enriched with repressive H3K27me3, forming bivalent promoters [55]. We assessed the H3K27me3 abundance at bivalent promoters in control and mutant mESCs using ChIP-seq. Consistent with H3.3 enrichment at bivalent promoters (n=4117; H3K27me3 + H3K4me3, see Methods) (**Fig. 6a**), the H3K27me3 signal at these regions was reduced in K27A mESCs (**Fig. 6b**), accompanied by a modest increase in H3K27ac (**Fig. 6c** and **Fig. S8d**). Additionally, we observed a decrease in the H3K27me3 signal at bivalent promoters in K9A mESCs. Histone mass-spectrometry data also revealed a global reduction in H3K27me3 levels in K9A mESCs (**Fig. S2**).

**Figure 6:**
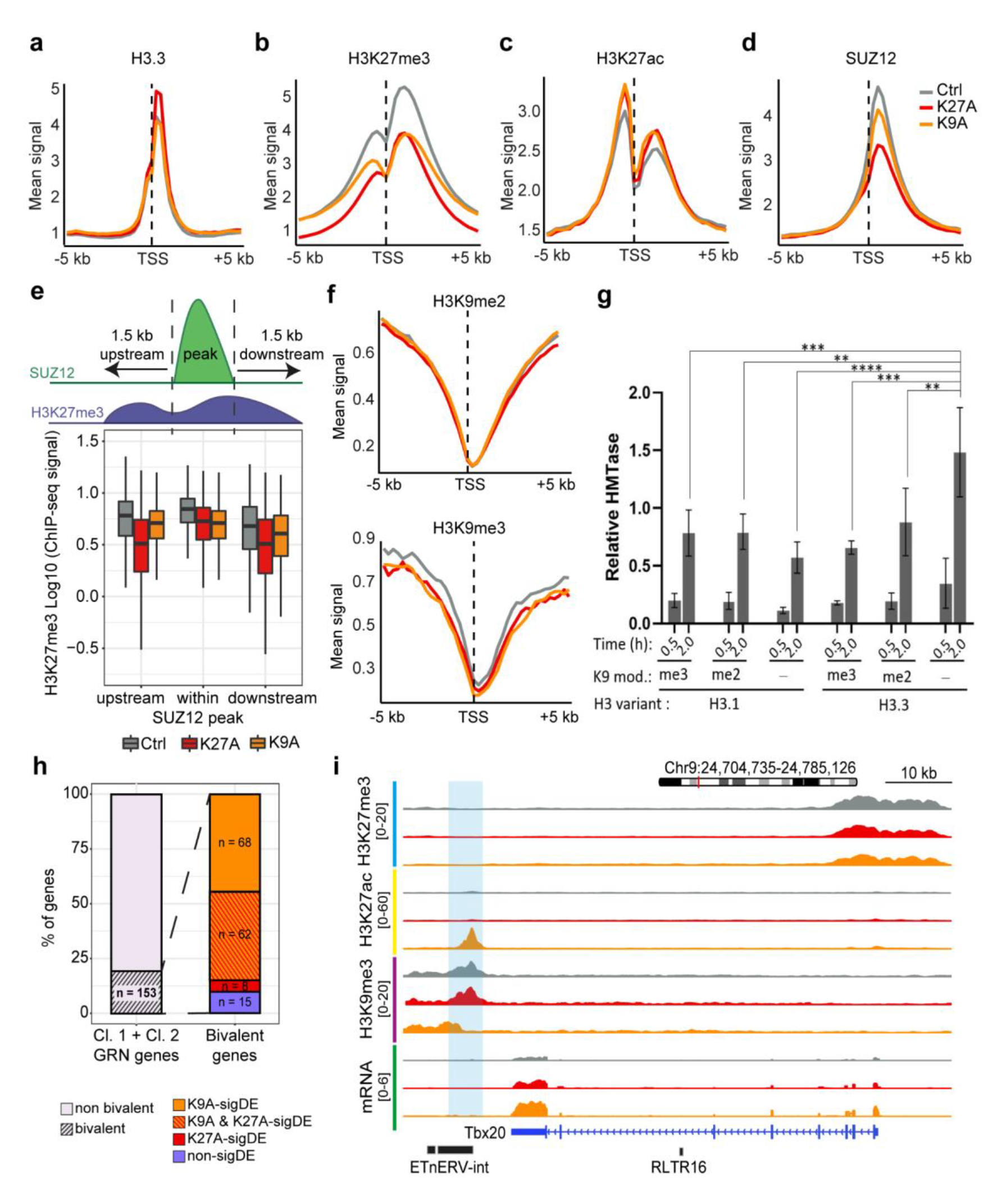
H3.3 K9A affected H3K27me3 level at bivalent promoters and increased bivalent genes via de-repressed cryptic enhancers. **(a-d)** Metaprofile plot of **(a)** H3.3, **(b)** H3K27me3, **(c)** H3K27ac, and **(d)** Suz12 ChIP-seq signal at TSS ± 5kb of bivalent genes (n=4117). **(e)** Analysis of H3K27me3 signal within SUZ12 peaks (n=3356) and in the 1.5 kb genomic region immediately upstream or downstream. **(f)** Metaprofile plot of H3K9me2 (left) and H3K9me3 (right) ChIP-seq signal at TSS ± 5kb of bivalent genes (n=4117). **(g)** Histone methyltransferase assays of PRC2 in the presence of chromatin reconstituted using different H3 variants and H3K9 modifications. In all cases, “me2” and “me3” represent the respective methyl lysine analog (MLA), and “-“ represents an unmodified lysine residue. Barplot represents the mean densitometry values as recorded from the radiograms (see Fig. S4c). The error bars represent the standard deviation over three replicates carried out on different days. *P* value: Tukey’s multiple comparisons test (****: P<0.0001, ***: P<0.001, **: P<0.005, *: P<0.05). **(h)** Stacked barplot indicating the percentage of genes with or without bivalent promoters (left), and details of significant differential expression (right). **(i)** Genome browser snapshot of a representative Cluster 2 DA-H3K9me3 region (highlighted), linked through the GRN to the Tbx20 gene; H3K27me3, H3K27ac, H3K9me3, and mRNA-seq tracks are shown. For metaprofiles in panels **a-d** and **f**, the ChIP-seq signal is calculated as the average of three biological replicates for control or mutant mESCs, in 250 bp sliding genomic windows.

We investigated the role of the H3.3 K9 residue in the proper deposition of H3K27me3 at bivalent promoters. As the PRC2 complex deposits H3K27me3, we mapped the occupancy of the PRC2 core subunit SUZ12 in control and mutant mESCs. SUZ12 binding was more substantially affected in K27A (1062 SUZ12 peaks showed a decrease) than K9A mESCs (41 peaks decreased: **Fig. 6d** and **Fig. S9a**), indicating a minimal effect of H3.3 K9 residue on PRC2 occupancy. Allosteric activation of PRC2 is required for the proper regulation of H3K27me3 deposition and spreading [56, 57]. We assessed the impact of the H3.3 K9A and K27A mutations on this process by analyzing H3K27me3 distribution over SUZ12 peaks [58]. While H3K27me3 signals within SUZ12 peaks were similarly reduced in both K27A and K9A mutants compared to control, H3K27me3 levels were further decreased in the immediate proximity of the SUZ12-bound regions in K27A mESCs only (**Fig. 6e**), indicating that the H3.3 K9 residue does not affect PRC2 allosteric activation.

Prior studies reported a putative function for H3K9 methylation in PRC2 activity, in which G9a and SETDB1 (H3K9 di-and tri-methyltransferases, respectively) reinforce PRC2 activity at target promoters [59–61]. ChIP-seq for H3K9me2 and H3K9me3 revealed that these marks were scarce at bivalent promoters (**Fig. 6f**). PRC2 activity assays using reconstituted chromatin composed of the ATOH1 bivalent locus DNA sequence combined with H3.1 or H3.3 histone variants marked with H3K9me2 or H3K9me3 did not show enhanced PRC2 activity (H3K27me3 increase; **Fig. 6g**). However, we found that chromatin reconstituted with un-methylated H3.3K9 was a better PRC2 substrate than H3.1K9 or di–/tri–methylated H3.3/H3.1K9 (**Fig. 6g**). This result suggests that the H3.3 K9 residue at Polycomb target loci could directly enhance the H3K27me3 signal, explaining the overall reduced H3K27me3 signal at PRC2-target promoters in K9A mESCs.

### De-repressed cryptic enhancers in H3.3K9A mESCs upregulate bivalent genes

Although the H3K27me3 signal was reduced at bivalent promoters, only a subset of bivalent genes showed a significant increase (fold-change cut-off of 2) in mutant mESCs. Bivalent genes upregulated in K27A and shared between the mutants (n=323) were enriched with typical developmental GO terms, whereas those upregulated only in K9A mESCs (n=340) showed divergent developmental GO terms such as synaptic transmission and leukocyte differentiation (**Fig. S9c**). We compared basal H3.3 occupancy at promoter regions of these genes using control mESCs. Bivalent genes upregulated in both mutants had significantly higher basal levels of H3.3 than those upregulated only in the K9A mutant or of randomly selected non-differential bivalent genes (**Fig. S9d**). This indicates that bivalent promoters with higher basal levels of H3.3 were more susceptible to the decrease in H3K27me3 upon H3.3K27A and H3.3K9A substitutions (i.e., H3.3K9 increases H3K27me3) and more likely to be de-repressed, suggesting direct causation. The cell lineage marker genes upregulated in both K27A and K9A mESCs (**Fig. 2a** and **Fig. S9c**) are examples of these bivalent promoter-containing genes.

We investigated whether the increased expression of bivalent genes exclusive to K9A mESCs resulted from the activation of distal regulatory elements. Among the genes included in the GRN, we identified 153 out of 789 DA-H3K9me3-linked genes with a bivalent promoter signature (**Fig. 6h**). Of those, 68 bivalent genes were exclusively upregulated in K9A mESCs, indicating that the de-repressed cryptic CREs could activate this subset of bivalent genes. Some of these genes, such as Tbx20 and Frzb (**Fig. 6i** and **Fig. S9e**), were involved in developmental processes and contained ERVs in the de-repressed CREs. Thus, we conclude that the upregulation of certain developmental genes in K9A mESCs is driven by the de-repression of cryptic ERV-derived CREs, similar to genes related to immune responses.

## Discussion

Here, our comparative systemic analysis demonstrates that H3.3K9 and H3.3K27 mediated PTMs are crucial for orchestrating repressive chromatin states at cis-regulatory elements and bivalent promoters, respectively, and for instructing proper transcription in mESCs. We have detailed how the removal of the K9 and K27 residues of histone H3.3 perturbs the histone modification landscapes in mESCs, and how those chromatin changes interplay to regulate gene expression.

Acetylation and methylation of H3K9 and H3K27 are involved in gene activation and repression, respectively. We found that changes in H3K9ac and H3K27ac in mutant mESCs merely correspond with enhancer activity. Our study revealed that a reduction in the level of H3K9me3 in H3.3K9A mESCs and H3K27me3 in H3.3K27A mESCs directly contributes to the gene activation observed in the individual mutants. In K9A mESCs, a decrease in H3K9me3 at numerous heterochromatic regions resulted in the activation of cryptic enhancers. Some of them have a distinctive genomic feature (i.e., Cluster 1) marked by H3K9me3, yet (i) harboring H3K27ac and (ii) having a higher enrichment for binding of TFs and transcriptional co-activators, compared to other heterochromatic regions in WT mESCs. These unique distal genomic regions, displaying high H3.3 occupancy, are therefore primed to be activated. We showed how these regions are strictly controlled by H3.3K9me3, and the reduction of this mark is sufficient to trigger a switch to an active chromatin state. By inferring enhancer-gene connections with the GRN generated in this study, we could identify genes presumably regulated by these cryptic CREs, such as those involved in immune and developmental processes.

We showed that the cryptic CREs activated in H3.3K9A mESCs derived from de-repressed ERV sequences. While histone H3.3 has been connected with the silencing of specific transposable elements, especially Class I and II ERVs, the contribution of H3.3K9 to ERV silencing has remained unclear because previous studies employed a H3.3 knockout approach [36, 62–65]. Given that we observed the de-repression of ERVs in mESCs solely by introducing the K9A substitution in the endogenous H3.3, we demonstrate that H3.3K9 is crucial for the silencing of ERVs.

Transposable element sequences contain binding sites for several TFs [39, 41, 66, 67]. The activation of ERV-derived CREs in H3.3K9A mESCs potentially creates competition with most active canonical enhancers (namely “strong”), by making TFs binding sites accessible in heterochromatic territories [54] and thus acting as sponges. Therefore, changes in H3K27ac occur indirectly in H3.3K9A mESCs and are overall correlated with rewired enhancer activities.

On the contrary, deposition of the H3K27ac mark is directly affected at canonical enhancers in H3.3K27A mESCs, and the H3K27ac decrease was accompanied by reduced acetylation at other H3 lysines. Nascent transcription levels were reduced, and the genes connected to these regions through the GRN were downregulated, suggesting that the overall level, rather than residue-specific, H3 acetylation may play a role in maintaining enhancers in an active state.

The modification status of histone residues can affect the deposition of other target residues [2, 68]. Here, our biochemical assay showed that unmodified H3.3K9 is a better substrate for PRC2 than H3.1K9, and thus the presence of H3.3K9A at Polycomb target promoters may have a direct negative impact on PRC2 activity, resulting in decreased H3K27me3 levels. Regarding the H3.3K27A effect on H3K27me3, consistent with recent studies in Drosophila [69] and mammals [17, 70–73], the H3.3K27 residue contributes to properly establishing the H3K27me3 mark at PRC2-bound promoters, as well as to the spreading of the modification. A decrease in promoter H3K27me3 levels does not always lead to gene activation. By comparing the bivalent genes upregulated in either or both mutants, we could identify two subsets: (i) one more susceptible to the decreases in the promoter H3K27me3 levels (comprising lineage marker genes) and affected in both mutants, and (ii) a subset less sensitive to H3K27me3 decrease, yet relying gene expression on the activation of distal regulatory elements and upregulated exclusively in H3.3K9A mESCs, due to the activation of ERV-derived CREs.

Notably, we observed in H3.3K9A mESCs the unusual activation of regulatory networks controlling the expression of immune genes, which also culminates in the expression of proteins typically expressed only in specialized immune cells. Although de-repressed ERVs may produce endogenous viral transcripts, which could trigger an interferon-mediated antiviral response, we did not observe a consistent and robust upregulation of interferon stimulated genes. This is in line with the reported limited ability of mESCs to mount innate immune responses. Together with the fact that ERVs have been co-opted throughout evolution to function as CREs in immune cells and other differentiated cell types, we envision that the ERV-derived cryptic CREs controlled by H3.3K9me3 in mESCs directly regulate the transcription of specific immune genes and developmental genes subsets, placing histone H3.3 lysine 9 as a critical caretaker of distinct distal genomic regions.

## Methods

### Cell culture

Mouse embryonic stem cells (129XC57BL/6J) were cultured in ESC medium containing Knockout-DMEM (Thermo Fisher Scientific) supplemented with 15% EmbryoMax fetal bovine serum (ES-FBS Merck-Millipore) and 20 ng/ml leukemia inhibitory factor (LIF, produced by Protein Expression and Purification Facility at EMBL Heidelberg), 1% non-essential amino acids (Thermo Fisher Scientific), 1% Glutamax (Thermo Fisher Scientific), 1% Pen/Strep (Thermo Fisher Scientific), 1% of 55 mM beta-Mercaptoethanol solution (Merck Millipore). Upon thawing, cells were cultured on a layer of mouse embryonic fibroblasts (MEFs) and afterwards passaged on 0.1% gelatine-coated culture dishes, without feeder cells. Cells were maintained at 37°C with 5% CO_2_ and routinely tested for mycoplasma. The medium was changed daily, and cells were passed every two days.

### CRISPR-Cas9 editing strategy

To introduce mutations of the endogenous *H3f3b* gene, a ribonucleoparticle (RNP) Cas9-based approach was used. First, tracrRNA-ATTO550 and the target specific crRNA-XT (Integrated DNA Technologies) were resuspended in IDT Nuclease-Free Duplex Buffer to a final concentration of 200 μM. The single-guide RNA (sgRNA; designed using the Benchling Guide RNA Design tool) was assembled *in-vitro* by mixing tracrRNA and crRNA in equimolar ratio, and the mixture was heated at 95°C for 5 min and cooled at RT on bench-top for ∼20 minutes. Second, the Cas9 RNP was assembled by mixing the Cas9-mSA protein (produced by the EMBL Protein Expression and Purification Facility) with the sgRNA in a molar ratio of 1:1.5 Cas9:sgRNA, in a final volume of 10 µl of Neon Buffer R per electroporation reaction and incubated at RT for 10-20 minutes. The biotinylated repair template (Integrated DNA Technologies Ultramers) was then added to the mix prior to electroporation, considering the same molar ratio of 1:1.5. Cells were washed once with PBS and 10^5^ cells/reaction were resuspended in 10 µl of mix previously prepared (Buffer R + Cas9 RNP complex). The Neon Transfection System (Thermo Fisher Scientific) was used for electroporation using the following program: 1600 V; 10 ms; 3 pulses (from Liang *et al.*, 2015 [74]). After electroporation, cells were transferred to a tube containing 500 μl of pre-warmed RPMI medium supplemented with 8% ES FBS and left to recover for 15 minutes at 37°C in the incubator. Cells were then centrifuged and washed once with RPMI medium supplemented with 8% ES FBS and plated in a 24-well plate containing MEFs. After 24 hours, single-cell sorting was performed to select ATTO550 positive cells and single clones were expanded, genotyped and validated. The nucleotide sequences of gRNA and repair templates used for the editing are reported in Table S1.

### Chromosomal integrity check of CRISPR-edited clones

The chromosomal integrity of the homozygous clones generated was assessed via low-coverage whole-genome sequencing. Briefly, genomic DNA was extracted from ∼10^6^ cells using the Gentra Puregene Cell Kit (Qiagen) and sonicated at 4°C using a Bioruptor Pico sonication device (Diagenode) for 10 cycles (30s ON/30s OFF) to obtain fragments of ∼200 bp. The fragmentation pattern was assessed via agarose gel electrophoresis and 0.5-1 μg of fragmented gDNA was used to prepare sequencing libraries using the NEBNext Ultra II DNA Library Preparation Kit (New England Biolabs), according to manufacturer’s instructions. Libraries were then quantified using the Qubit fluorometer (Thermo Fisher Scientific) and the quality was assessed on a Bioanalyzer using the Agilent High Sensitivity DNA Kit. Samples were then pooled and sequenced on a Illumina NextSeq-500 platform (75 bp single-end reads).

### Low-coverage whole-genome sequencing analysis

Quality of the sequencing run was assessed using FastQC (v 0.11.5 [75]) and adapter sequences were trimmed using TrimGalore (v 0.4.3 – https://github.com/FelixKrueger/TrimGalore). Sequencing reads were aligned to the mouse reference genome (GRCm38/mm10 assembly) using Bowtie2 (v 2.3.4 [76]) with default settings. Uniquely mapped reads (MAPQ ≥ 30) were retained for the subsequent steps. The bam files thus generated were then indexed and used for copy number variation analysis (bin size of 100 kb) using the Coral script (https://github.com/tobiasrausch/coral). Pre-processing steps were conducted with the Galaxy platform (v 21.09 [77]).

### Histones extraction and mass-spectrometry analysis

Histones were extracted, derivatized with propionic anhydride, and digested as previously described [78]. All samples were desalted prior to nanoLC-MS/MS analysis using in-house prepared C18 stage-tips. Histone samples were analyzed by nanoLC-MS/MS with a Dionex-nanoLC coupled to an Orbitrap Fusion mass spectrometer (Thermo Fisher Scientific). The column was packed in-house using reverse-phase 75 µm ID × 17 cm Reprosil-Pur C18-AQ (3 µm; Dr. Maisch GmbH). The HPLC gradient was: 4% solvent B (A = 0.1% formic acid; B = 80% acetonitrile, 0.1% formic acid) for 2 minutes, then from 4% to 34% solvent B over 48 min, then from 34% to 90% solvent B in 2 minutes, hold at 90% solvent B for 2 minutes, and from 90% solvent B down to 4% in 1 minute followed by a hold at 4% solvent B for 5 minutes. The flow rate was at 300 nL/min. Data were acquired using a data-independent acquisition method, consisting of a full scan MS spectrum (m/z 300−1200) performed in the Orbitrap at 120,000 resolution with an AGC target value of 5e5, followed by 16 MS/MS windows of 50 m/z using HCD fragmentation and detection in the orbitrap. HCD collision energy was set to 27, and AGC target at 1e4. Histone samples were resuspended in buffer A and 1 ug of total histones was injected. Histone data was analyzed using a combination of EpiProfile 2.0 [79], Skyline [80], and manual analysis with Xcalibur (Thermo Fisher Scientific). The peptide relative ratio was calculated by using the area under the curve (AUC) for that particular peptide over the total AUC of all possible modified forms of that peptide. Data analysis was performed using Microsoft Excel to calculate averages and standard deviations.

### PRC2 *in-vitro* HMT assay

The assay was performed as previously described [81], with the addition of normalization samples that were loaded on each of the gels to allow for a comparison across multiple gels. Each 10 μL HMTase reaction contained 500 nM PRC2 complex, chromatinized DNA with sequence from the ATOH1 locus (see Zhang *et al.*, 2021 [82]) at a total concentration equivalent to 0.8 μM nucleosomes and 5.0 μM 14C-SAM (S- [methyl-14C]-adenosyl-l-methionine (PerkinElmer, no. NEC363050UC)). All chromatinized DNA constructs were identical except for the H3 histone constructs, which included either H3.1 or H3.3, as indicated, and lysine 9 was either unmodified or modified to include a methyl lysine analog (MLA), as indicated. Each reaction was incubated for 30 minutes and 2 h at 30 °C in HMTase buffer (50 mM Tris-HCl pH 8.0 (at 30°C), 100 mM KCl, 2.5 mM MgCl2, 0.1 mM ZnCl2, 2.0 mM 2-mercaptoethanol and 0.1 mg/mL-1 bovine serum albumin) and then stopped by adding 4× LDS sample buffer (∼3.3μL) (Thermo Fisher Scientific, no. NP0007) to a final concentration of 1× (final volume ∼13.3μL). 10 μL of each sample was then incubated at 95 °C for 5 min and subjected to SDS-PAGE (16.5% acrylamide). Gels were first stained with InstantBlue Coomassie protein stain (Expedeon, no. ISB1L) and then vacuum-dried for 60 min at 80 °C with the aid of a VE-11 electric aspirator pump (Jeio Tech). Dried gels were exposed to a storage phosphor screen (Cytiva) for 5 days and the signal was acquired using a Typhoon Trio imager (Cytiva). All experiments were performed in triplicate.

### Growth assay

mESCs were seeded at a density of 10^4^ cells per well and I seeded 12 wells/line on day 0. Two wells per cell line were trypsinized and counted each day for a total of 6 days, while the remaining wells received fresh media. I generated growth curves using the averaged duplicate cell counts.

### Cell cycle assay

The percentage of proliferating cells in S-Phase was measured using the EdU Click FC-488 Kit (Carl Roth – BA-7779) following the manufacturer’s instructions. Briefly, EdU was added at a final concentration of 10 µM and cells were incubated for 2.5 hours at 37°C. Cells were washed once with PBS and then dissociated using accutase. Cells were fixed by resuspension in 100 µl (every 1×10^6^) of 4% PFA in PBS and incubation 15 minutes at RT, in the dark. After quenching with PBS + 1% BSA, cells were centrifuged and the pellet resuspended in 1X saponin-based permeabilization buffer. A total of 4.5×10^5 cells were used for each condition and 500 µl of Click reaction master mix (PBS, catalyst solution, dye azide and 10X buffer additive) were added to each tube. After 30 minutes incubation at RT in the dark, cells were washed once with 3 ml of saponin-based permeabilization buffer and DNA content was stained with propidium iodide (7 minutes incubation at RT). Cells were washed once with saponin-based permeabilization buffer and resuspended in 300-500 µl of buffer and analyzed by flow cytometry. Three controls were included for flow cytometry analysis: i) w/o EdU, w/o click and w/o DAPI; ii) w/o Edu, with click and DAPI; iii) with EdU, w/o click and with DAPI.

### Cell death assay

The FITC Annexin V Kit (BD Pharmingen – 556419) was used, which allows to detect earlier stages of cell death, preceding loss of membrane integrity. Briefly, cells were washed twice with PBS and resuspended in 1X binding buffer to a concentration of 1×10^6^ cells/ml. To 100 µl of the mix (∼ 10^5^ cells), 5 µl of FITC Annexin V and 10 µl of propidium iodide (50 µg/ml stock) were added and cells were incubated for 15 minutes at RT, in the dark. After incubation, 400 µl of 1X binding buffer were added to each tube and cells were analyzed by flow cytometry. Three controls were included for flow cytometry analysis: i) unstained cells; ii) cells with FITC Annexin V, w/o PI; iii) cells w/o FITC-Annexin V, with PI.

### Immune proteins staining

Cells were harvested and washed with FACS buffer (PBS + 2% FBS). Cells were then resuspended in 100 µl of master mix composed of FACS buffer and desired antibodies, each one diluted 1:100. Alternatively, cells were resuspended in FACS buffer only for the unstained controls. After 30 minutes incubation on ice and in the dark, cells were centrifuged and washed with FACS buffer. Antibodies used for the staining were the following: PE-CD59a (143103 – Biolegend); APC-TLR2/CD282 (153005 – Biolegend); FITC-MHC Class II (107605 – Biolegend).

### Bodipy-C11 staining

The staining was performed following the guidelines described by Martinez *et al.*, 2020 [83]. In brief, the BODIPY-C11 probe was added at a final concentration of 2.5 µM and cells were incubated for 30 minutes at 37°C. Cells were then washed with HBSS buffer, collected and analyzed by flow cytometry.

### Neuronal differentiation

mESCs were differentiated into glutamatergic neurons, as described by Bibel *et al.*, 2007 [84], with modifications. mESCs were cultured in differentiation medium containing high-glucose DMEM, 10% FBS (Gibco), 1% non-essential amino acids (Thermo Fisher Scientific), 1% penicillin/streptomycin (Thermo Fisher Scientific), 1% GlutaMAX, 1% sodium pyruvate (Thermo Fisher Scientific), 0,1% of 14,5 M β- mercaptoethanol (Sigma Aldrich) solution. To promote the formation of embryoid bodies, cells were grown in suspension on non-adherent dishes (Greiner Petri dishes – 633102). The differentiation medium was changed every two days. From day 4 until day 8, the EBs were treated with 5 μM retinoic acid every two days to induce neuronal lineage commitment. On day 8, EBs were dissociated with trypsin and neural progenitor cells (NPCs) were plated on poly-D-lysine hydrobromide (Sigma Aldrich) and laminin (Roche) coated dishes in N2 medium (high-glucose DMEM, 1% N-2, 2% B-27 and 1% penicillin/streptomycin, from Thermo Fisher Scientific) at a density of 2×10^5^ cells/cm^2^. The medium was changed after 2 h and after 24 h. On day 10, the medium was changed to complete medium (neurobasal, 2% B-27 and 1% penicillin/streptomycin). Glutamatergic neurons were harvested on day 12.

### Mesoderm-cardiomyocyte differentiation

mESCs were differentiated into cardiomyocytes as previously described [18], adapting a protocol from previous publications [85, 86]. Briefly, 7.5×10^5^ mESCs were resuspended in differentiation medium (DMEM, 20% FBS, 1% NEAA, 1% P/S, 1% GlutaMAX, 100 μM ascorbic acid) and grown in suspension on non-adherent dishes to promote EB formation. After 4 days, EB were plated onto 0,1% gelatine-coated plates. The medium was changed every two days. Formation of contracting colonies was observed after 8 days of differentiation and time-course quantification of contracting colonies was performed at day 8, 10 and 12. Cardiomyocytes were harvested by trypsinization on day 14.

### Quantitative real-time PCR

RNA was extracted from approximately 10^6^ cells using RNeasy Kit (Qiagen), followed by DNase digestion using TURBO DNase (Thermo Fisher Scientific). A total of 1 μg of RNA was reverse transcribed with random primers to cDNA using High-Capacity cDNA Reverse Transcription Kit (Applied Biosystems). For qRT-PCR reaction, 9 ng of cDNA were used as template and reactions were performed using Power SYBR Green master mix (Applied Biosystems) on a StepOne Plus Real-Time PCR machine. The comparative Ct method (ΔΔCt method) was used to calculate normalized gene expression values. Ct values of target genes were normalized to Ct values of the housekeeping gene Rpl13 to obtain ΔCt values and to control samples to obtain ΔΔCt values. Primers used for qRT-qPCR of NPCs markers are listed in the Table S2.

### Immunofluorescence

NPCs were seeded on glass coverslips at day 8 of neuronal differentiation and fixed at day 12 for neuronal staining. Cells were washed with PBS briefly and fixed with 3% paraformaldehyde (PFA) (Electron Microscopy Sciences) in PBS for 20 minutes at RT, then quenched with 30 mM glycine in PBS for 5 minutes at RT. Cells were washed three times with PBS and stored in PBS at 4°C until needed. Permeabilization was conducted with 0.1% Triton-X in PBS for 5 minutes at RT, followed by blocking with 0.5% BSA in PBS for 30 minutes at RT. Primary antibody incubation was performed for 1 hour at RT under constant shaking. The following antibodies were diluted 1:200 in 0.5% BSA and used for stainings: Map2 (Sigma Aldrich, 9942) and β-III tubulin (Abcam, ab78078). Cells were washed three times with PBS and incubated for 30 minutes at RT with secondary antibody (goat anti-mouse IgG Alexa 594, Thermo Fisher Scientific – A11005), diluted 1:1000 in 0.5% BSA. After washing twice with PBS, cells were counterstained with 5 μg/ml DAPI for 5 minutes at RT and washed three times with PBS. Coverslips were mounted with Mowiol mounting medium (Calbiochem) and imaged on a Nikon Eclipse Ti fluorescence microscope. Images were processed with Fiji ImageJ.

### mRNA-seq

RNA was extracted from ∼10^6^ cells using the RNeasy Mini kit (Qiagen) and DNase digestion was performed using TURBO DNase (Thermo Fisher Scientific), according to the manufacturer’s instructions and an additional clean-up of the RNA was then performed using the RNeasy Mini kit (Qiagen). Quality of the RNA was assessed on Bioanalyzer using the Agilent RNA 6000 Nano Kit and samples with a RNA integrity number (RIN) of 10 were used for library generation. For each sample, 100 ng of total RNA was used as input for mRNA selection and conversion to cDNA using the NEBNext Poly(A) Magnetic Isolation Module (New England BioLabs). Sequencing libraries were prepared using the NEBNext Ultra II RNA Library Preparation Kit for Illumina (New England BioLabs), according to the manufacturer’s instructions. After Qubit quantification, the quality of individual libraries was assessed on a Bioanalyzer using the Agilent High Sensitivity DNA Kit. Samples were then pooled and sequenced on an Illumina NextSeq500 sequencer (read length of 75 bp in single-end mode).

### mRNA-seq analysis

Quality of the sequencing reads was assessed using FastQC (v0.11.5) and adapter sequences were trimmed using TrimGalore (v0.4.3 - https://github.com/FelixKrueger/TrimGalore). Sequencing reads were aligned to the mouse reference genome (GRCm38/mm10 assembly) using STAR (v2.5.2b [87]) with default settings. Uniquely mapped reads (MAPQ ≥ 20) were retained for the subsequent steps. Gene count tables were generated with featureCounts (subread v1.6.2 [88]), using gencode gene annotations (release M10). Coverage files were generated using bamCoverage (deeptools v2.4.1 [89]). Differential expression analysis was performed in R using the DESeq2 package [90]. Genes were considered differentially expressed using a false discovery rate (FDR) cut-off of 0.05 and the fold-change cut-off applied is indicated in the main text and in figure captions. MA-plots were generated using the ggmaplot function (from the ggpubr R package – v0.4.0; https://github.com/kassambara/ggpubr). Upset plot in Figure 2.5 was generated using the UpSetR package [91]. Gene ontology enrichment analysis was conducted using ClusterProfiler [92]. Other plots were generated using the ggplot2 R package (https://github.com/tidyverse/ggplot2). Pre-processing steps were conducted with the Galaxy platform (v21.09).

### Transposable elements expression analysis

For endogenous retroviruses (ERVs) expression analysis, the RepEnrich tool [93] was used in its updated version (https://github.com/nerettilab/RepEnrich2). Sequencing reads were aligned to the mouse reference genome (GRCm38/mm10 assembly) using Bowtie2 (v2.3.4), and retaining secondary alignments. LTRs annotation for the mm10 mouse genome assembly was retrieved using the RepeatMasker track from the UCSC genome table browser and prepared as suggested. After running RepEnrich2 independently for all samples, the individual output tables with estimated counts were merged, and the differential expression analysis was performed in R using DESeq2. Repeats were considered differentially expressed using a FDR cut-off = 0.05 and abs(log2 fold-change) cut-off = 0.585. The PCA plot was generated in R with ggplot2 and volcano plots were generated with the EnhancedVolcano package (https://github.com/kevinblighe/EnhancedVolcano). For the overlap analyses (Fig. 5a) and for inspection of specific loci on genome browser (Figg. 5h, 6i, S7a and S9e), only ERVs with size ≥500 bp were considered.

### Native ChIP-seq

DNA for native ChIP was digested by MNase treatment to obtain mainly mono-nucleosomes using a modified protocol from Barski *et al.*, 2007 [94]. For each IP, 20×10^6^ cells were resuspended in digestion buffer (50 mM Tris-HCl pH 7.6; 1 mM CaCl_2_; 0.2% Triton-X), treated with 100 U MNase (Worthington) and incubated at 37°C for 5 minutes while shaking at 500 rpm. Samples were quickly moved to ice and MNase was quenched by addition of 5 mM EDTA (final). Lysates were sonicated for 3 cycles using Bioruptor Pico (Diagenode) and dialyzed against RIPA buffer for 3 hours at 4°C. Insoluble materials was pelleted at 10,000 rpm for 10 minutes at 4°C and supernatant was used as input for ChIP. To check the fragmentation pattern, input DNA was analyzed by agarose gel electrophoresis. A 5% fraction of input was set aside for sequencing. Protein-G-Dynabeads (Invitrogen) were pre-coated with antibodies for 4 hours at 4°C and the following antibodies were used: H3K27ac (39685 – Active Motif); H3K9ac (C5B11-9549 – Cell Signalling Technology). After overnight incubation of the pre-coated beads with chromatin lysates, beads were washed (3X with RIPA; 2X with RIPA + 300 mM NaCl; 2X with LiCl buffer–250 mM LiCl, 0.5% NP-40, 0.5% NaDoc) and finally rinsed with TE + 50 mM NaCl Buffer. Samples were eluted from ProteinG Dynabeads using SDS elution buffer (50 mM Tris-HCl ph 8.0; 10 mM EDTA; 1% SDS) at 65°C for 30 minutes with shaking at 1500 rpm. Proteinase K was added (0.2 µg/ml final) to the eluted samples and digestion was carried out for 2 hours at 55°C followed by PCR purification (Qiagen). Sequencing libraries were prepared using the NEBNext Ultra II DNA Library Preparation Kit (New England Biolabs) and sequenced on Illumina NextSeq500 platform (75 bp in single-end mode).

### Cross-link ChIP-seq

The protocol was adapted from Groh *et al.*, 2021 [38] and Navarro *et al.*, 2020 [37]. Cells were harvested and crosslinked in 3 ml of pre-tempered (25°C) ES medium supplemented with 1% formaldehyde (10^6^ cells/3 ml) for 10 minutes at RT, with rotation. Formaldehyde was then quenched with glycine (125 mM final) and incubated 5 minutes at RT, with rotation. Cells were washed twice with ice-cold PBS containing 10%FBS (centrifugation at 1100 rpm for 4 minutes, at 4°C). Pellets can be stored at –80°C for several months. To prepare lysates, fixed cells were resuspended in 300 µl of Sonication Buffer 1 (50 mM Tris-HCl pH 8.0; 0.5% SDS) and incubated for 10 minutes on ice. Sonication was performed with Bioruptor Pico (Diagenode) for 20 cycles (30’’ON/30’’OFF). Sonicated lysates were then diluted 1:6 with lysis buffer (10 mM Tris-HCl ph 8.0; 100 mM NaCl; 1% Triton-X; 1 mM EDTA; 0.5 mM EGTA; 0.1% NaDoc; 0.5% N-laurolsarcosine) and the soluble fraction was collected after centrifugation at full speed for 10 minutes at 4°C. A 2.5% fraction of the supernatant was set aside for input and the fragmentation pattern (∼150–500 bp) was checked by agarose gel electrophoresis. Lysates can be aliquoted and stored at –80°C with 10% glycerol (final). For each IP, 30 µl Protein G Dynabeads (Invitrogen) were washed twice with 1 ml PBS-T (PBS + 0.01% Tween-20) and incubated with the desired antibody for at least 1 hour at RT (or > 1 hour at 4°C). Antibodies used with this protocol were the following: H3.3 (09-838 – Merck-Millipore); H3K9me3 (D4W1U – Cell Signalling Technology); H3K9me2 (D85B4 – Cell Signalling Technology); H3K27me3 (C36B11 – Cell Signalling Technology); H3K18ac (9675 – Cell Signalling Technology); SUZ12 (D39F6-3737 – Cell Signalling Technology).

Coated beads were then washed (1X with PBS-T; 2X with lysis buffer), resuspended in 30 µl of lysis buffer/IP and added to the desired amount of chromatin (generally 15-25 µg of chromatin were used, corresponding to 2-4×10^6^ cells). After overnight incubation at 4°C with rotation, beads-immunocomplexes were washed twice—each time for 5 minutes—with the following buffers: RIPA; RIPA + 360 mM NaCl; LiCl buffer (10 mM Tris-HCl ph 8.0; 250 mM LiCl, 0.5% NP-40, 0.5% NaDoc; 1 mM EDTA) and finally quickly rinsed with TE buffer and eluted in ChIP SDS elution buffer (10 mM Tris-HCl ph 8.0; 300 mM NaCl; 5 mM EDTA; 0.5% SDS). RNA/protein digestion was performed on beads by adding 2 µl RNaseA (10 mg/ml stock) incubated 30 minutes at 37°C, followed by addition of 1.5 µl Proteinase K (20 mg/ml stock) incubated 1h @ 55°C. Reverse cross-link was performed with overnight incubation at 65°C. DNA was purified with 1.4X SPRI-select beads. For H3K27me3 ChIP-Rx, spike-in chromatin was prepared from Drosophila Schneider 2 (S2) cells following the same protocol and aliquots were stored at –80°C with 10% glycerol. Before immunoprecipitation, 3% of exogenous chromatin was added to each reaction. Sequencing libraries were prepared using the NEBNext Ultra II DNA Library Preparation Kit (New England Biolabs) and sequenced on Illumina NextSeq500 (75 bp in single-end mode) or NextSeq2000 (P3 kit – 88 bp in single-end mode).

### ChIP-seq analysis

Quality of the sequencing run was assessed using FastQC (v0.11.5) and adapter sequences were trimmed using TrimGalore (v0.4.3 - https://github.com/FelixKrueger/TrimGalore). Sequencing reads were aligned to the mouse reference genome (GRCm38/mm10 assembly) using Bowtie2 (v2.3.4). Uniquely mapped reads (MAPQ ≥ 20) aligning to major chromosomes were retained for the subsequent steps. ChIP signal strength and sequencing depth were assessed using the plotFingerprint and plotCoverage (deeptools v2.4.1). Coverage files (bigwig format) were generated using deeptools bamCoverage: 10 bp was used as bin size, the “reads per genomic content (RPGC)” method was used for normalization and reads were extended to an average fragment size of 150 bp. For the H3K27me3 ChIP-seq with exogenous spike-in, reads were aligned to a mouse (mm10) + Drosophila (dm6) combined reference genome. The number of uniquely mapped reads aligning to major mouse and fly chromosomes was retrieved and used to calculate normalization factors that were used with deeptools bamCoverage to generate scaled bigwig files. Peak calling was performed with MACS2 [95], by providing ChIP and respective input bam files and considering a minimum FDR cut-off for peak detection of 0.05; narrow or broad peak calling was performed for histone modifications following ENCODE guidelines (https://www.encodeproject.org/chip-seq/histone/). Differential peak analysis was performed in R using the DiffBind package [96], only peaks detected in at least two replicates were retained for the analysis and peaks were called as significantly differential considering an FDR cut-off = 0.05. Annotation of ChIP-seq peaks was performed with the ChIPseeker package [97] and with publicly available ChromHMM state maps for mESCs (https://github.com/guifengwei/ChromHMM_mESC_mm10).

Annotations for bivalent promoters were obtained using previously generated H3K4me3 and H3K27me3 ChIP-seq data [18]. Briefly, a consensus peakset was defined for the control mESCs (peaks identified in at least two replicates) for both datasets; regions of overlap for the two histone marks were obtained using findOverlaps-methods and genomic regions within a 500 bp window were merged. Genomic coordinates of promoters of protein coding genes were retrieved from ENSEMBL using biomart (mm10 version: https://nov2020.archive.ensembl.org) and only H3K4me3-H3K27me3 regions overlapping with TSS ± 1 kb were retained. This annotation was further refined by selecting promoters overlapping with peaks of PRC2 subunits (i.e., SUZ12; EZH2; EZH1; EED) retrieved from the ChIP-Atlas database [24] and finally validated with ChromHMM state maps for mESCs.

Heatmaps and metagene plots were generated in R: genomic regions of interest were binned in 40 windows of 250 bp each, the ChIP-seq signal from individual bigwig files was measured in each window and all bigwig regions overlapping the same window were averaged. The signal for replicates in each condition was then averaged. Heatmaps were generated using the pheatmap package (https://cran.r-project.org/web/packages/pheatmap/index.html). Boxplots and violin plots were generated considering the ChIP-seq signal measured at TSS or peak summit ± 1 kb. Overlaps between genomic regions of interest were quantified using the GenomicRanges objects and findOverlaps-methods.

Genome browser shots in Figg. 5h, 6i, S7a and S9e were generated using the Gviz package [98]; bigwig files used for the visualization represent the average ChIP-seq signal from the three biological replicates of each genotype, which was computed using WiggleTools [99].

### ChIP-Atlas data integration

Publicly available ChIP-seq data for chromatin factors and transcription factors in mESCs (mm10 assembly) were retrieved from the ChIP-Atlas database [24] and the analysis was performed with an approach similar to the one previously described in Bunina *et al.*, 2020 [100]. Briefly, genomic coordinates of peaks for all mESCs factors (n=292) were retrieved from the ChIP-Atlas Peak Browser and the findOverlaps function was used to overlap these peaks with genomic regions of interest. The significance of the relative enrichment of chromatin/transcription factors was assessed using Fisher’s exact test. Factors were selected as follows: (i) significant in DA-regions versus non-DA regions (randomly selected H3K9me3 and H3K27ac genomic regions considered as background), (ii) significant in each cluster versus other clusters; only factors significant in both comparisons were retained and used for STRING network analysis.

### Transposable elements ChIP-seq analysis

Analysis of H3.3, H3K9me3 and H3K27ac ChIP-seq data to detect enrichment at transposable elements was carried out using the Transposable Element Enrichment Estimator (T3E) tool [101]. Sequencing reads were aligned to the mouse reference genome (GRCm38/mm10 assembly) using Bowtie2 (v2.3.4), retaining secondary alignments. T3E was run for all the ChIP replicates and input samples, filtering out genomic regions of high signals (as suggested by the developers for mouse genome). A total of 20 iterations were performed when running T3E and calculating enrichments. ChIP-seq enrichment at TEs was considered significant with a p-value<0.01 and ERVs with high H3.3 occupancy were additionally filtered by applying a log2(fold-change)>1 cut-off.

### ChIP-qPCR

Prepared libraries from ChIP experiments were diluted and 5 ng of DNA was used as input for each qPCR reaction with SYBR Green PCR Master Mix (Applied Biosystems). The qPCR reactions were performed on a StepOnePlus Real-Time PCR machine. For each condition biological replicates (ChIP material from two independent mutant cell lines) were used and measured in technical duplicates. Primers used for H3K9me3 and H3K18ac ChIP-qPCR are reported in Table S2.

### PRO-seq

Precision nuclear run-on transcription sequencing (PRO-seq) experiments were performed following the protocol previously described [102]. PRO-seq was performed in two batches, one with two replicates each of control and H3.3K27A mESCs and one with three replicates each of control and H3.3K9A mESCs. Briefly, 10^7^ mESCs were permeabilized and used for each reaction and 5% (i.e., 500k) of permeabilized Drosophila S2 cells were added prior to the *in-vitro* run-on reaction. Run-on reactions were carried out by adding 100 µl of 2× NRO reaction mixture (10 mM of Tris-HCl, pH 8.0; 5 mM MgCl_2_; 300 mM KCl; 1 mM dithiothreitol (DTT); 1% sarkosyl; 375 µM biotin-11-CTP/-UTP; 375 µM ATP/GTP; 2 µl RNase inhibitor) to 100 µl of the permeabilized cells, and incubated at 30 °C for 3 min. Addition of TRIzol LS (Thermo Fisher Scientific) terminated the reaction and RNA was extracted and precipitated in 75% ethanol. Nuclear RNA was fragmented by base hydrolysis in 0.2 N of NaOH on ice for 10 min and subsequently neutralized by adding the same volume of 1 M of Tris-HCl, pH 6.8. Fragmented biotinylated RNA was bound to the streptavidin magnetic beads (Thermo Fisher Scientific), washed, eluted using TRIzol and precipitated in ethanol. Following 3′–RNA and 5′–RNA adapter ligation, the precipitated RNA was reverse-transcribed and PCR-amplified. DNA libraries were purified using SPRI-select beads and quality was assessed on a Bioanalyzer using the Agilent High Sensitivity DNA Kit, pooled and sequenced on an Illumina NextSeq 500 platform (75 bp in single-end mode).

### PRO-seq analysis

The analysis of the PRO-seq data was performed according to the guidelines provided by the Danko Laboratory. In particular, pre-processing and alignment was performed using the PROseq2.0 pipeline (https://github.com/Danko-Lab/proseq2.0). The consensus set of distal *cis*-regulatory elements active at the basal state in mESCs was obtained by running the dREG package (https://github.com/Danko-Lab/dREG) on the PRO-seq data from control cell lines. The differential PRO-seq analysis was performed using the output of the dREG package from control and H3.3K9A lines as input for the tfTarget package (https://github.com/Danko-Lab/tfTarget). The violin plots in Figg. 3e, 5i and S8f were generated by calculating RPKM values for mutant and control samples within the genomic regions of interest.

### Gene regulatory network construction and differential transcription factor activity analysis

The GRN was generated as described in Kamal & Arnold *et al.*, 2022 [25], integrating mRNA-seq and H3K27ac ChIP-seq datasets obtained from 12 independent mESCs samples (i.e., three biological replicates each for Control, H3.3K9A, H3.3K27A and H3.3K79A mESCs respectively) which were used to construct the GRN. An FDR cutoff=0.2 was used for peak-gene connections, yielding a GRN with TF-CRE-gene connections composed of a network of 153 unique TFs, 5694 regulatory elements, 5229 genes (of which 1194 DE) overall resulting in 16701 CRE-gene connections.

The differential transcription factor activity analysis was performed using the diffTF method described in Berest & Arnold *et al.*, 2019 [29], using the same 12 mRNA-seq and ChIP-seq datasets as input.

## Acknowledgments

We thank the staff of the European Molecular Biology Laboratory (EMBL) Genomics Core Facility, Protein Expression and Purification Facility, and Flow Cytometry Core Facility for sample preparations and data generation. We thank Charles Girardot and the Genome Biology Computational Support for their assistance in data analysis and submission. We thank Mohamad Whaidi, Maja Gehre, and Nichole Diaz for helping with mutant lines generation. We thank Andrea Callegari and Moritz Kueblbeck for sharing their expertise in CRISPR-Cas9 gene editing. We thank Michael Bonadonna for helping with flow-cytometry stainings. We thank Brian Mang Ching Lai for helping with the GRN construction. We thank Elena Vizcaya Molina for providing the Drosophila S2 cells used as spike-in in PRO-seq and ChIP-seq experiments. We thank Guy Riddihough for critical feedback on this manuscript and Eileen Furlong and the Noh laboratory for the helpful discussions.

## Author contributions

M.T. and K.M.N. conceived the project. M.T. and K.M.N. designed the experiments. M.T., D.B., U.Y., N.F., and A.J. collected and analyzed the data. M.T. and D.B. performed the bioinformatic analyses. J.Z. supervised D.B. M.U., V.L., and C.D. designed and performed the PRC2 in-vitro assays. Y.K. performed the histone modifications mass-spectrometry analysis with supervision from B.A.G.. M.T. and K.M.N. wrote the manuscript with input from all authors.

## Supplementary figures

**Figure S1:**
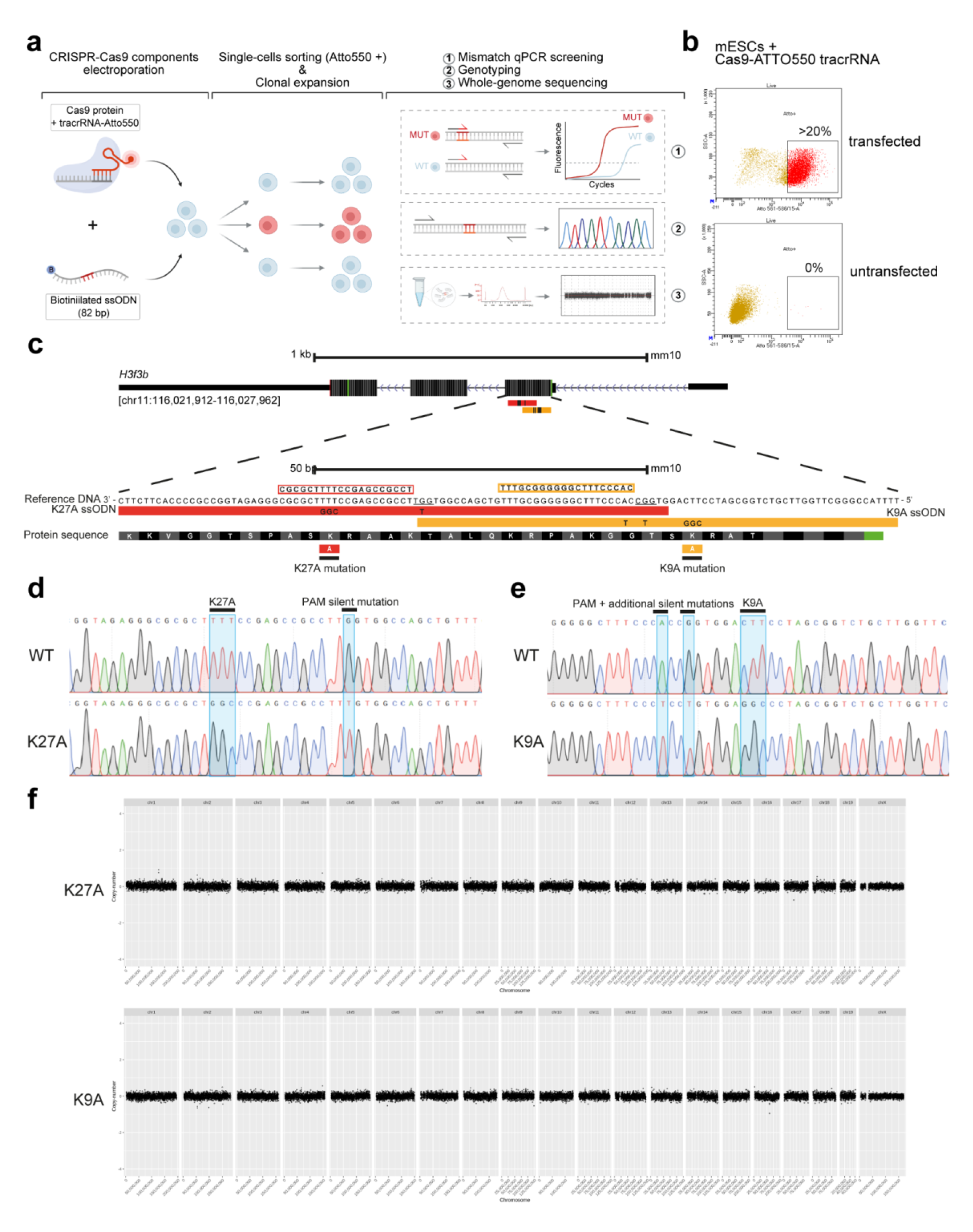
CRISPR-Cas9 editing strategy and chromosomal integrity assessment for the edited clones. **(a)** Schematic describing the Cas9 RNP editing workflow (created with BioRender.com) **(b)** Flow cytometry analysis displays mESCs transfection efficiency with Cas9-tracrRNA-ATTO550 (about 20% or higher - top). Untransfected cells subjected to Cas9 RNP complex but not transfected are used to set the gates for flow cytometry analysis (bottom). **(c)** Schematic with guide RNAs and ssODN designed to introduce K27A (red) and K9A (orange) mutations in the *H3f3b* locus. **(d-e)** Representative Sanger sequencing chromatograms showing the comparison between wild-type and K27A (d) or K9A (e) mESCs at the *H3f3b* endogenous locus. **(f)** Representative whole-genome plots showing the copy-number difference of 100 kb genomic bins between edited clones and the parental control line.

**Figure S2:**
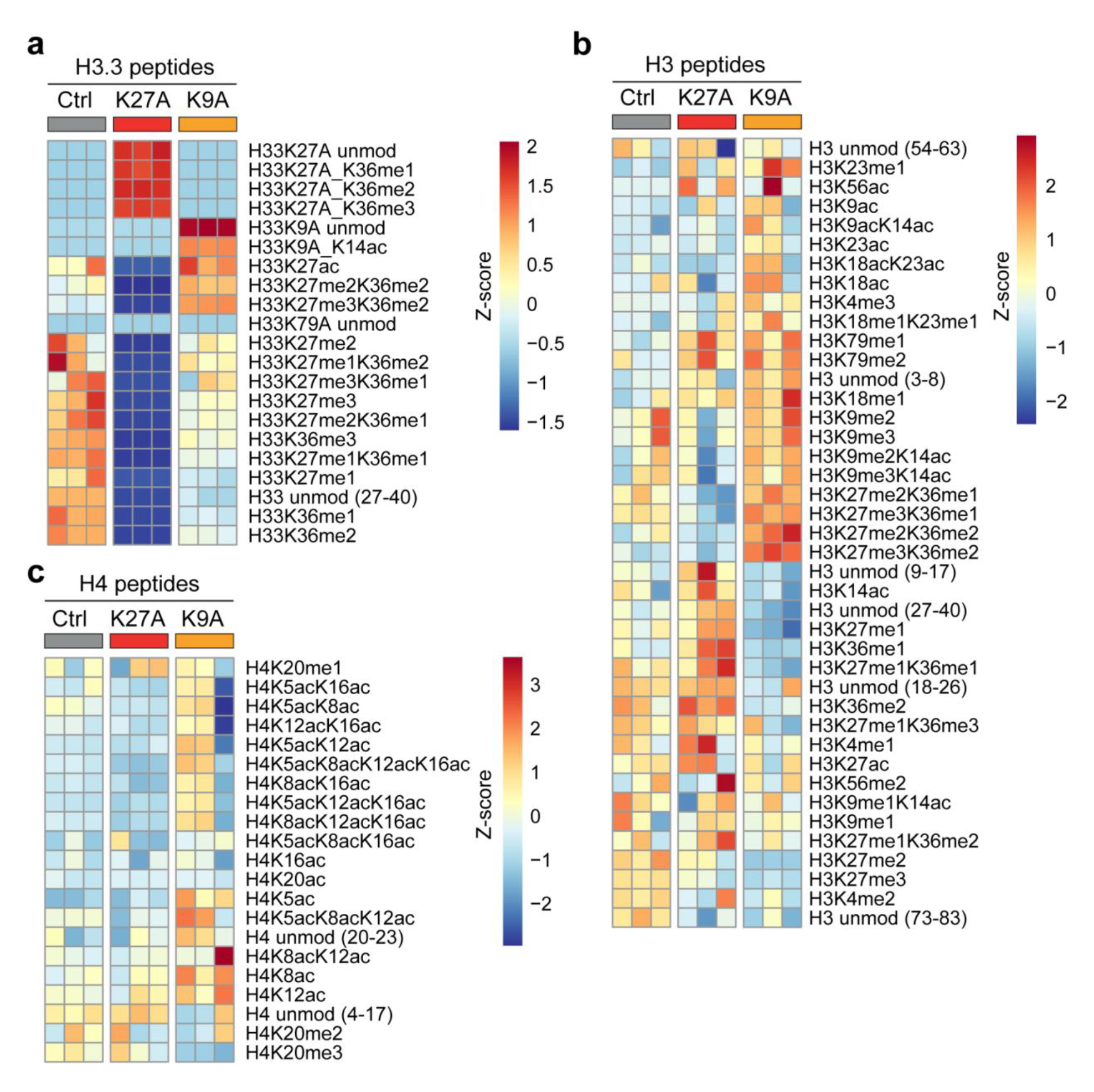
Histone modifications middle-down mass spectrometry analysis. Heatmaps displaying the relative abundance (row z-score) of individual or combinatorial post-translational modifications of histone H3.3 **(a)**, H3 **(b)** or H4 **(c)**. Lysine-to-alanine mutated H3.3 peptides were used as controls in panel “a”. The peptides length was omitted for clarity from the figure and is reported here: H3K4 (3-8); H3K9/K14 (9-17); H3K18/K23 (18-26); H3K27/K36 (27-40); H3K56 (54-63); H3K79 (73-83); H4K20 (20-23); H4K5/K8/K12/K16 (4-17); H4K20 (20-23).

**Figure S3:**
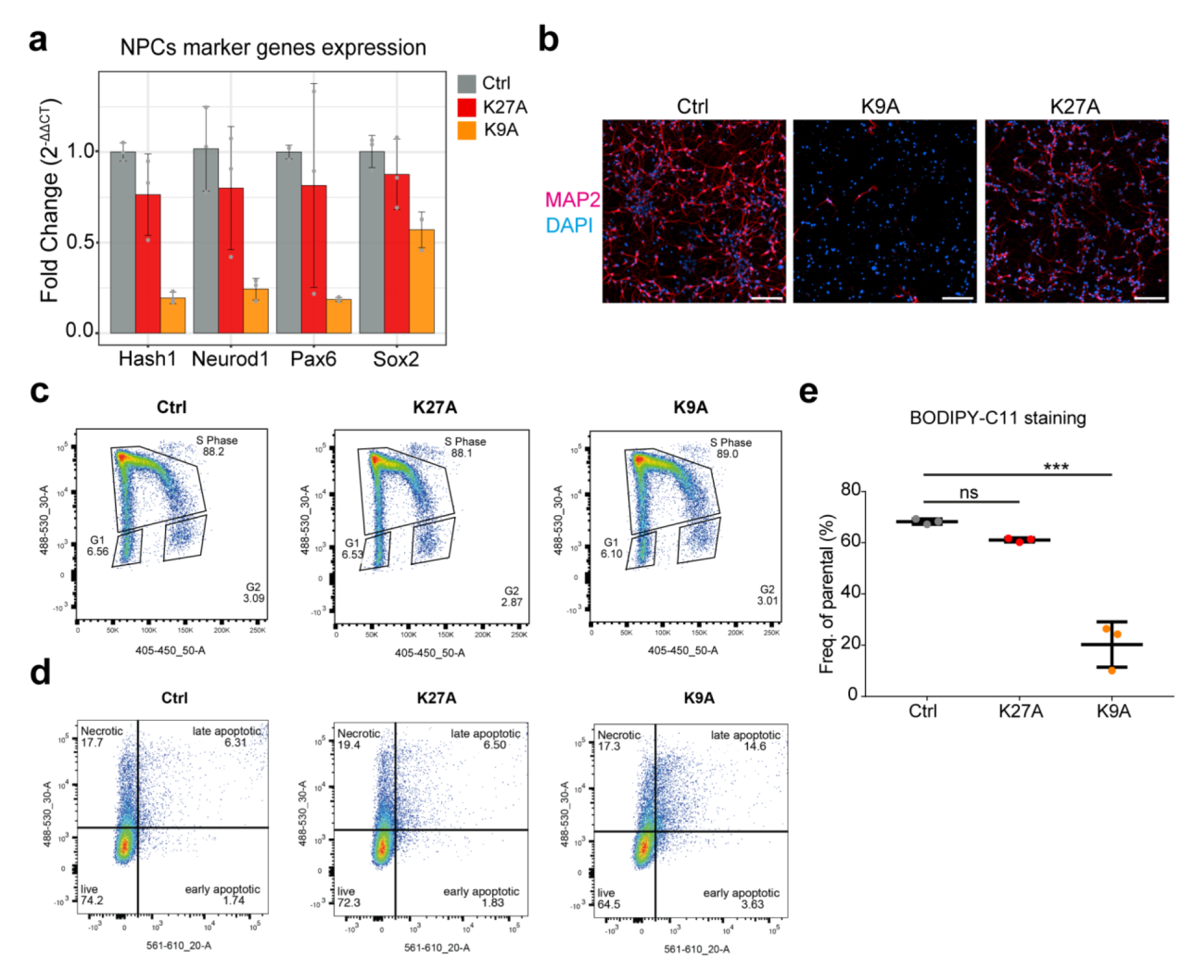
Phenotypic characterization of H3.3 K27A and K9A mESCs. **(a)** qRT-PCR results for neuronal progenitor cell markers (day 8) in control, K9A and K27A (n=3 independent clonal lines/condition). Data are the mean ± standard deviation of gene expression values (2−ΔΔCt) normalized to Rpl13 housekeeping gene and control. **(b)** Merged immunofluorescence images of neurons on day 12 of the *in-vitro* differentiation, stained with antibodies against MAP2 and with DAPI to detect nuclei. Scale bar 100 μm. **(c)** Representative plots of flow cytometry analysis displaying gating strategy for measuring 5-ethynyl-2’-deoxyuridine (EdU) incorporation in control, K27A and K9A mESCs. **(d)** Representative plots of flow cytometry analysis upon Annexin V staining for control, K27A and K9A mESCs. **(e)** Levels of lipid peroxides measured in control, K27A and K9A mESCs using BODIPY-C11 staining and flow-cytometry analysis. Significance was calculated with unpaired t-test (***: p-value<0.001; ns: p-value>0.05).

**Figure S4:**
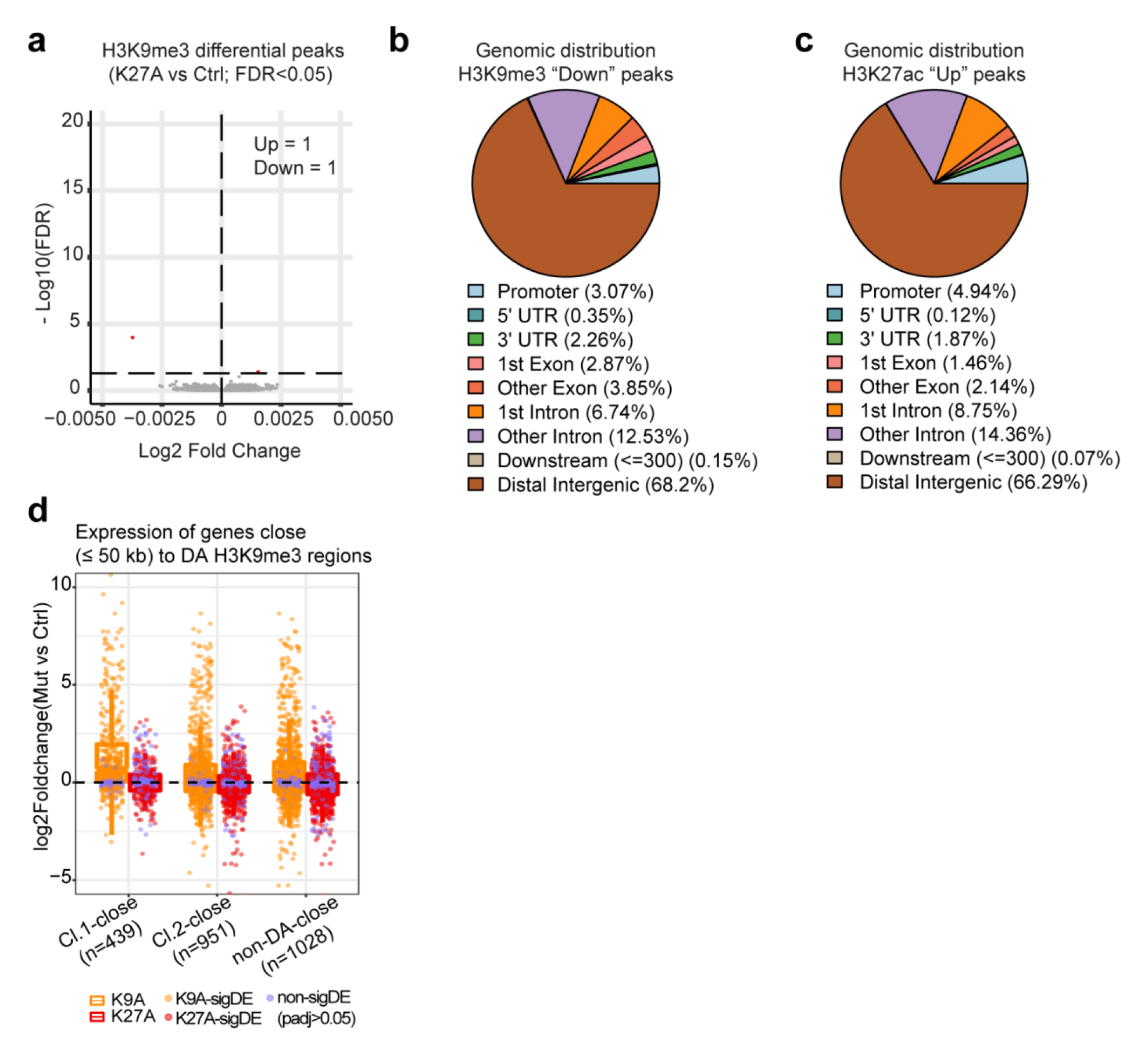
Genomic distribution of H3K9me3-lost and H3K27ac-gained regions. **(a)** Volcano plots showing differential H3K9me3 ChIP-seq analysis in K27A. **(b)** Genomic annotation of the H3K9me3 regions with significantly reduced signal in K9A mESCs. Promoter defined as TSS ± 1 kb. **(c)** Genomic annotation of the H3K27ac regions with significantly increased signal in K9A mESCs. Promoter defined as TSS ± 1 kb. **(d)** Expression of genes located within 50 kb from DA-H3K9me3 regions; individual genes (data points) are coloured to indicate if the differential expression is significant (orange/red) or not (purple).

**Figure S5:**
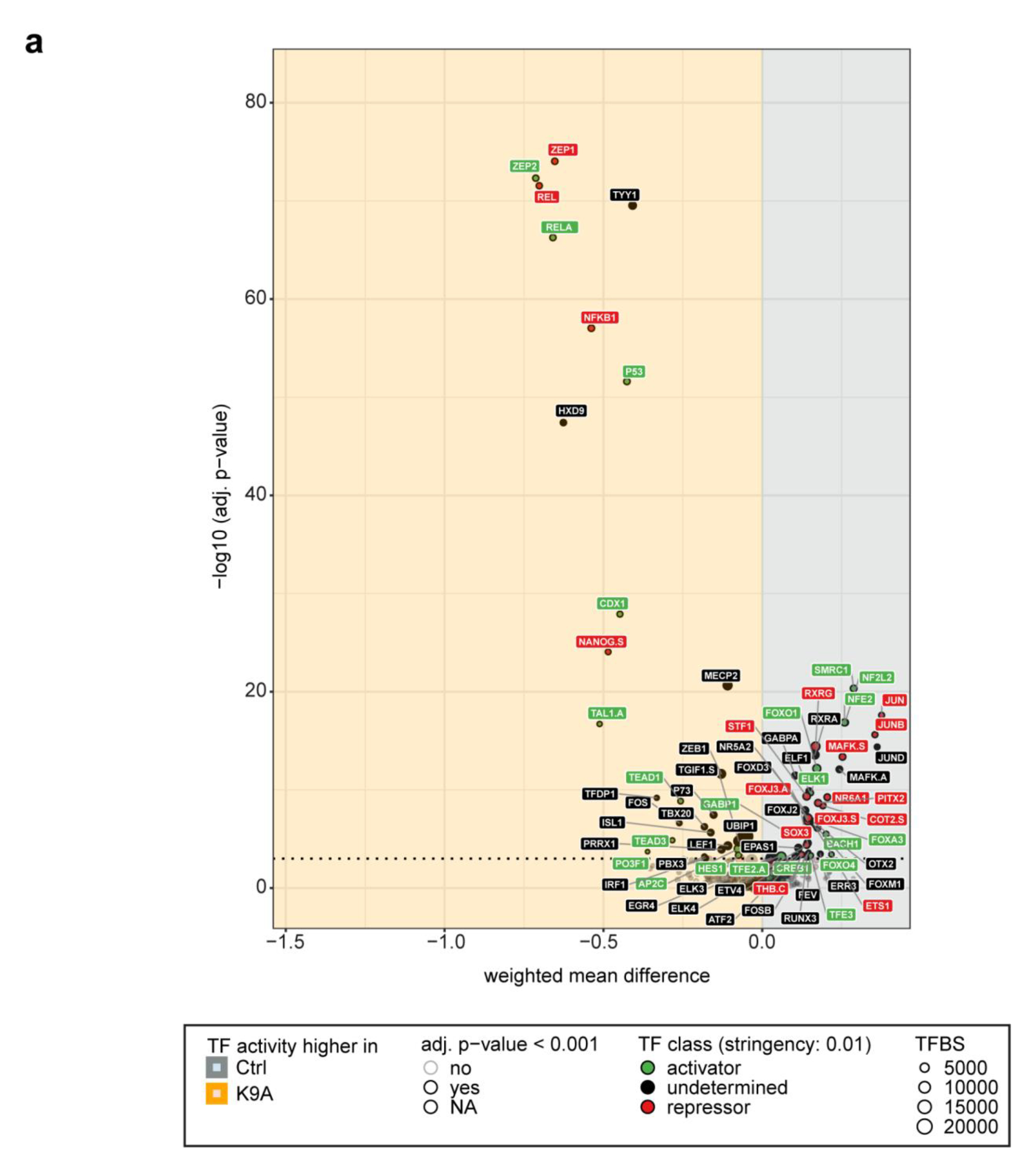
diffTF analysis for K9A mESCs. **(a)** Volcano plot showing TFs with higher activity in K9A (left – orange) or in control (right – gray). TFs are classified as activators (green) or repressors (red). Only expressed TFs are displayed.

**Figure S6:**
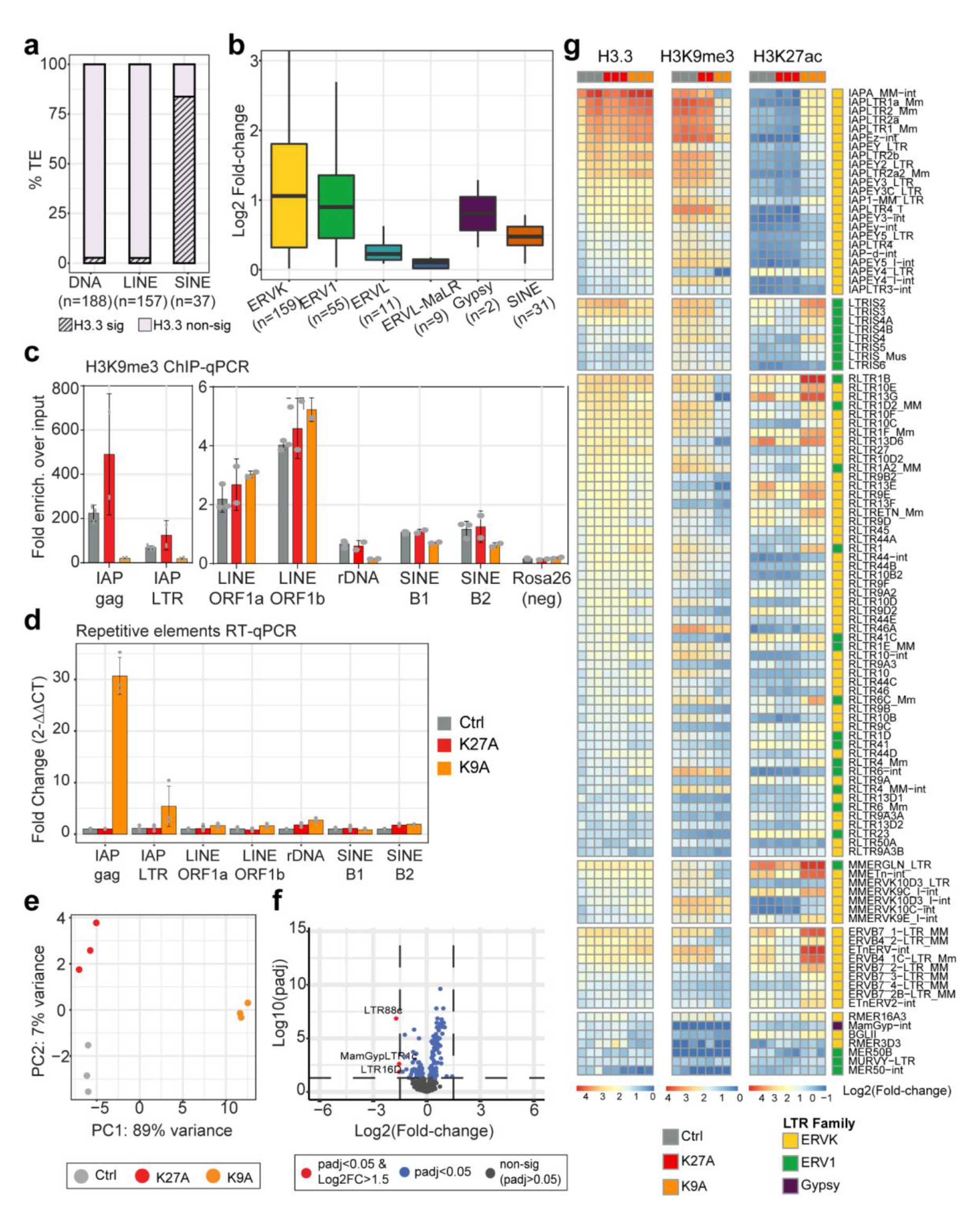
TE analyses. **(a)** Stacked barplot indicating the percentage of transposable elements with or without significant (pval<0.01) H3.3 enrichment over input in control mESCs. **(b)** Boxplot of H3.3 log2(Fold-change) enrichment at ERVs families and SINE elements, displaying significant H3.3 signal (from panel a and Fig. 5b) **(c)** H3K9me3 ChIP-qPCR with primers for ERVs (left) or other transposable elements (right). Rosa26 locus was used as a negative control for H3K9me3 enrichment. **(d)** Real-time qPCR using primers for selected transposable elements. **(e)** Principal component analysis of LTRs expression. **(f)** Volcano plot showing differential ERVs expression in K27A mESCs. **(g)** Heatmaps displaying H3.3, H3K9me3 and H3K27ac log2(Fold-change) enrichment at ERVs subfamilies with highest H3.3 signal (pval<0.01 & log2FC>1; n=108). ERVs subfamilies are divided in six groups, and sorted in each group by decreasing H3.3 signal in control mESCs. For each subfamily, the corresponding ERV family is reported and color-coded (ERVK in yellow; ERV1 in green and Gypsy in purple).

**Figure S7:**
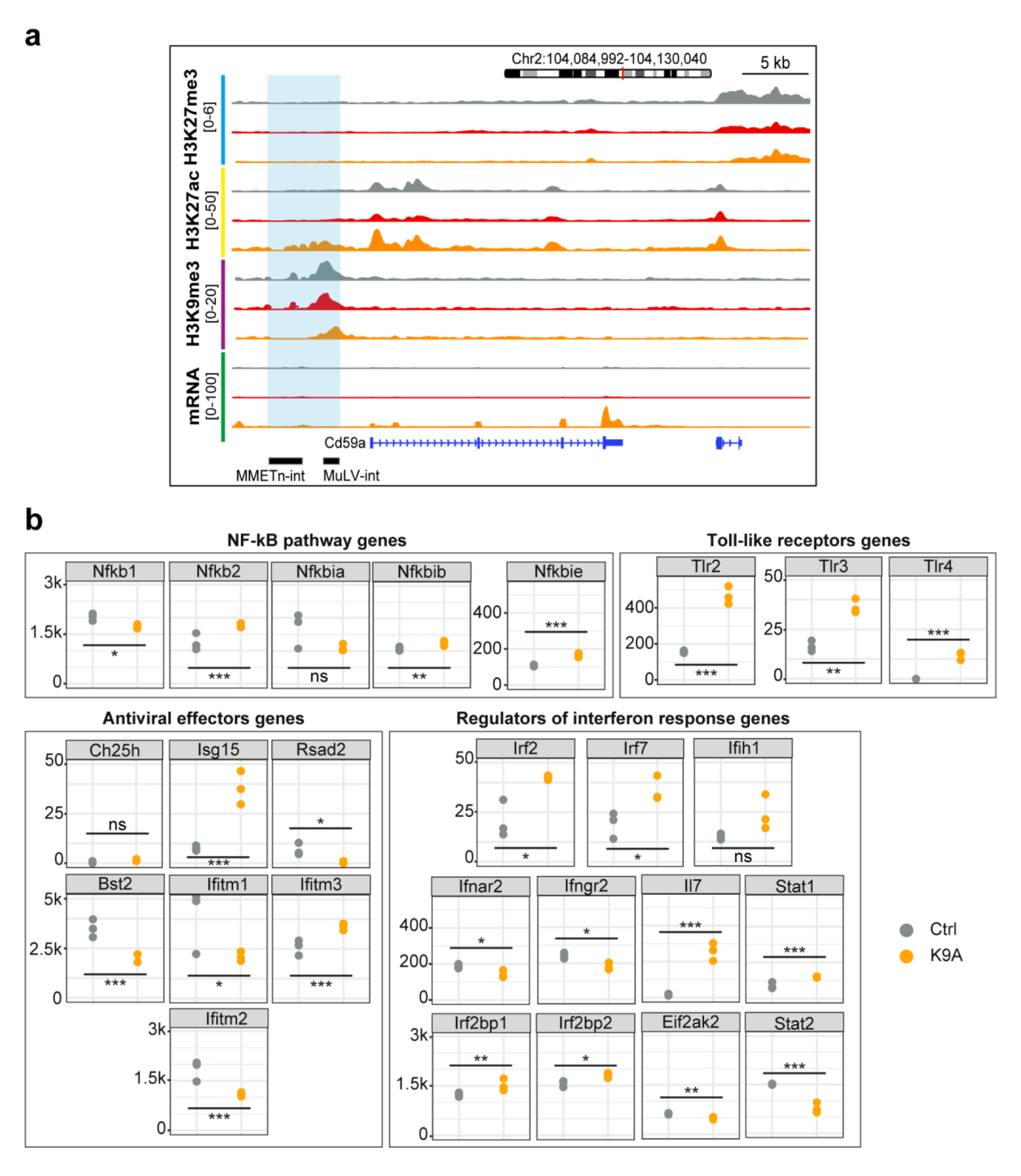
Cd59a genome browser shot and expression of interferon response genes. **(a)** Genome browser snapshot of a representative Cluster 2 DA-H3K9me3 region (highlighted), linked through the GRN to the Cd59a gene. From top to bottom, H3K27me3, H3K27ac, H3K9me3, and mRNA-seq tracks are shown. **(b)** Normalized mRNA-seq counts for selected interferon-stimulated genes in control and K9A mESCs. Genes are divided in four groups: NF-kB pathway genes (Nfkb1/p50; Nfkb2/p65; Nfkbia/IkB-alpha; Nfkbib/IkB-beta; Nfkbie/IkB-epsylon), Toll-like receptors genes, Antiviral effectors genes (Ch25h; Isg15; Rsad2/Viperin; Bst2/Tetherin; Ifitm1-2-3) and Regulators of interferon response genes (Irf2-7; Ifih1/Mda5; Ifnar2; Ifngr2; Il7; Irf2bp1-2; Eif2ak2/Pkr; Stat1-2). Significant differences were calculated with DESeq2 and adjusted with Benjamini Hochberg’s correction. (*: p-adj<0.05; **:p-adj<0.01; ***:p-adj<0.001; ns: p-adj>0.05).

**Figure S8:**
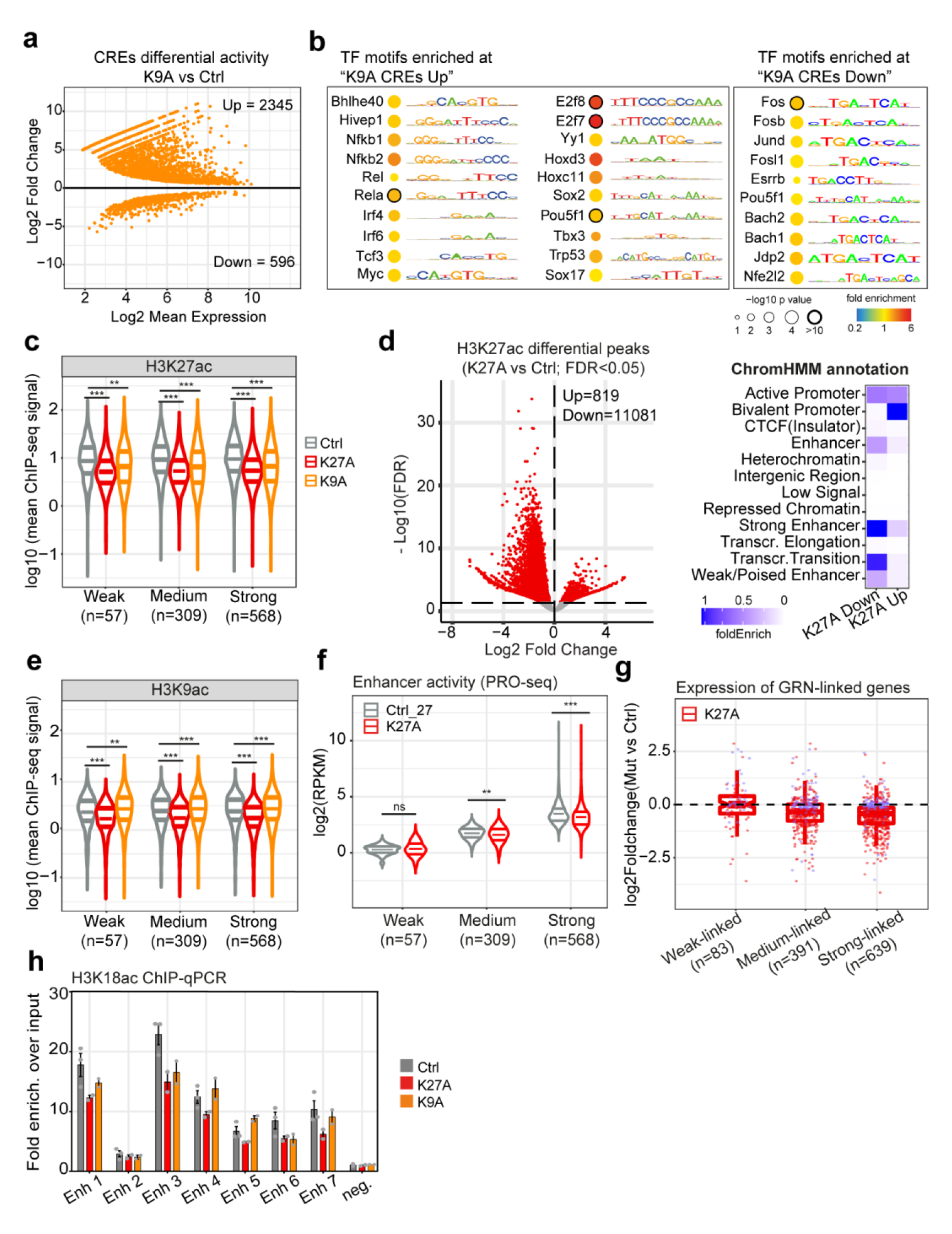
TF target analyses and H3K27ac and H3K9ac at canonical enhancers. **(a)** MA-plot showing differentially active CREs, according to tfTarget analysis (retained CREs as significant with padj<0.01). **(b)** Selected TF motifs significantly enriched at CREs with increased (left) or decreased (right) activity in K9A mESCs. **(c)** Violin plots showing H3K27ac ChIP-seq signal at peak center ± 1 kb of enhancers retained in the GRN. **(d) (e)** Violin plots showing H3K9ac ChIP-seq signal at peak center ± 1 kb of enhancers retained in the GRN. **(f)** PRO-seq signal at Weak, Medium, and Strong dREG enhancers included in the GRN for K27A and control mESCs. Log2(RPKM) values are reported. **(g)** Expression of DEGs connected to dREG enhancers through the GRN; log2(fold-change) values calculated with DESeq2 are plotted, and individual genes (data points) are colored to indicate if the differential expression is significant (red) or not (purple). In panels “c”, “d” and “f”, significance was calculated with unpaired t-test (*: p-value<0.05; **: p-value<0.01; ***: p-value<0.001, ns: p-value>0.05). **(h)** H3K18ac ChIP-qPCR at selected Strong enhancers regions. The relative enrichment over the input is reported and data are mean ± standard deviation (n=3 replicates Ctrl; n=2 replicates mutants).

**Figure S9:**
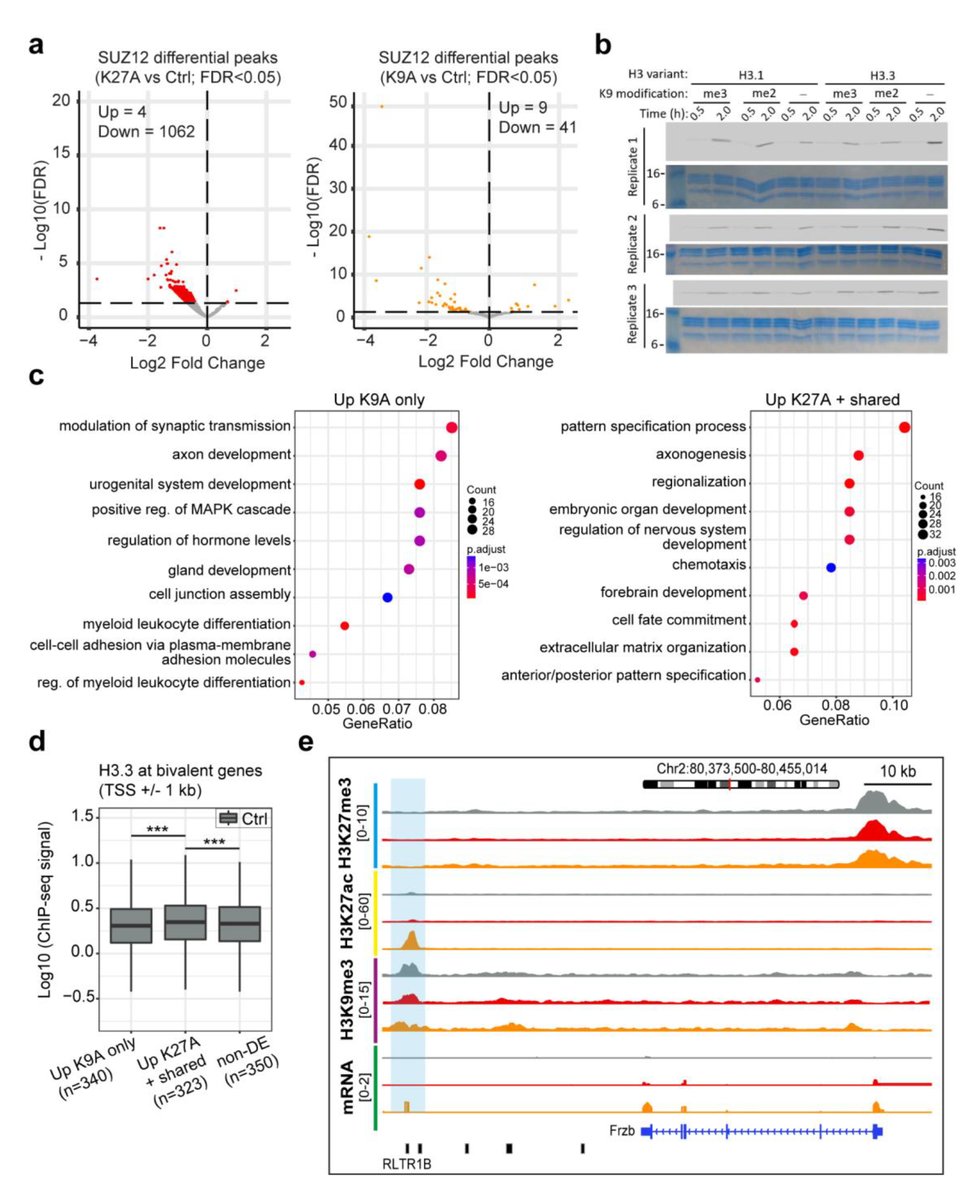
Effect of H3.3 K27A and K9A mutations on PRC2. **(a)** Volcano plots showing differential SUZ12 ChIP-seq peaks in K27A (left) or K9A (right) versus control (consensus peakset with n=3356 peaks). **(b)** Radiograms used to calculate mean values reported in Fig. 6g for PRC2 methyltransferase assay. **(c)** Gene ontology enrichment analysis of bivalent genes upregulated only in K9A (left) or in K27A/K27A+K9A mESCs (right). **(d)** H3.3 ChIP-seq signal in control mESCs, at the promoter regions of bivalent genes, upregulated only in K9A or K27A/K27A+K9A. A group of comparable size with non-significantly differentially expressed bivalent genes was randomly selected for comparison. Significance was calculated with unpaired t-test (***: p-value<0.001). **(e)** Genome browser snapshot of a representative Cluster 2 DA-H3K9me3 region (highlighted), linked through the GRN to the Frzb gene. For ERVs annotation, only the name of the ERV located within the DA-H3K9me3 region is shown; from left to right the other elements are: RMER10A, RLTR20B2, RLTR13B1 and RMER15-int.

